# Continuous Bump Attractor Networks Require Explicit Error Coding for Gain Recalibration

**DOI:** 10.1101/2024.02.12.579874

**Authors:** Gorkem Secer, James J. Knierim, Noah J. Cowan

**Affiliations:** Laboratory for Computational Sensing and Robotics, Johns Hopkins University, Baltimore, MD 21218, USA; Zanvyl Krieger Mind/Brain Institute, Johns Hopkins University, Baltimore, MD 21218, USA; Solomon H. Snyder Department of Neuroscience, Johns Hopkins University School of Medicine, Baltimore, MD 21205, USA; Kavli Neuroscience Discovery Institute, Johns Hopkins University School of Medicine, Baltimore, MD 21205, USA; Department of Mechanical Engineering, Johns Hopkins University, Baltimore, MD 21218, USA

## Abstract

Representations of continuous variables are crucial to create internal models of the external world. A prevailing model of how the brain maintains these representations is given by continuous bump attractor networks (CBANs) in a broad range of brain functions across different areas, such as spatial navigation in hippocampal/entorhinal circuits and working memory in prefrontal cortex. Through recurrent connections, a CBAN maintains a persistent activity bump, whose peak location can vary along a neural space, corresponding to different values of a continuous variable. To track the value of a continuous variable changing over time, a CBAN updates the location of its activity bump based on inputs that encode the changes in the continuous variable (e.g., movement velocity in the case of spatial navigation)—a process akin to mathematical integration. This integration process is not perfect and accumulates error over time. For error correction, CBANs can use additional inputs providing ground-truth information about the continuous variable’s correct value (e.g., visual landmarks for spatial navigation). These inputs enable the network dynamics to automatically correct any representation error. Recent experimental work on hippocampal place cells has shown that, beyond correcting errors, ground-truth inputs also fine-tune the gain of the integration process, a crucial factor that links the change in the continuous variable to the updating of the activity bump’s location. However, existing CBAN models lack this plasticity, offering no insights into the neural mechanisms and representations involved in the recalibration of the integration gain. In this paper, we explore this gap by using a ring attractor network, a specific type of CBAN, to model the experimental conditions that demonstrated gain recalibration in hippocampal place cells. Our analysis reveals the necessary conditions for neural mechanisms behind gain recalibration within a CBAN. Unlike error correction, which occurs through network dynamics based on ground-truth inputs, gain recalibration requires an additional neural signal that explicitly encodes the error in the network’s representation via a rate code. Finally, we propose a modified ring attractor network as an example CBAN model that verifies our theoretical findings. Combining an error-rate code with Hebbian synaptic plasticity, this model achieves recalibration of integration gain in a CBAN, ensuring accurate representation for continuous variables.

## 1 Introduction

The brain’s ability to represent and process continuous variables, such as location, time, and sensory information, is fundamental to our understanding and interaction with the external world. A compelling theoretical framework for how the brain constructs these representations is provided by continuous bump attractor networks (CBANs) in a diverse range of brain functions, such as orientation tuning in visual cortex [1], working memory [2,3], evidence accumulation and decision-making [4–7], and spatial navigation [8–11].

The CBAN is a class of recurrent neural network, which maintains persistent patterns of population activity through interactions among its neurons. This persistent activity typically forms the shape of a bell curve like a ‘bump’, when visualized on an appropriate topological arrangement of neurons (known as a low-dimensional manifold), such as a plane, circle, or torus [12]. Although the shape of the activity bump is constrained by network dynamics, its center location can vary along this low-dimensional manifold, corresponding to different values of the encoded continuous variable. Neural activity consistent with these key properties of CBANs, namely, the activity bump and the low-dimensional manifold, have been observed in recordings from various regions of the mammalian brain that encode continuous variables [13–15]. More conclusive and direct evidence for CBANs has been found in the central complex of fly brain, where a biological CBAN encoding the fly’s heading angle, a continuous variable, has been identified based on the connectome and a combination of techniques, such as calcium imaging and optogenetics [16–18]. While these experimental findings support the idea of brain circuits employing CBANs to represent continuous variables, the neural mechanisms that enable CBANs to accurately update their representations in response to changes in continuous variables remain incompletely understood.

CBANs update their representations of a continuous variable based on one or both of two distinct types of inputs. The first type provides ‘absolute’ information, namely, the true value of a continuous variable, such as spatial location relative to visual landmarks or the item to be held in the working memory. When this absolute information is available, it provides localized input on the CBAN’s low-dimensional manifold to a location that is associated with the true value to the continuous variable. In response to this localized input, internal dynamics of the CBAN creates a ‘basin’ on its low-dimensional manifold toward which the activity bump gravitates, resulting in nearly perfect encoding of the continuous variable [19–21]. Strong experimental evidence for this phenomenon has been observed in fly brain where optogenetic excitation of the central complex, a seemingly biological CBAN, at a specific anatomical location mimicking an absolute information resulted in the CBAN’s activity bump, normally changing its location to encode the fly’s heading, being pulled to the excitation location [16]. As more indirect evidence, neural recordings from the hippocampus and entorhinal cortex, two regions modeled as CBANs in mammalian brain, showed that their representations for the animal’s location are anchored to the visual landmarks even when the landmarks are rotated around an open arena [22–26].

In contrast to the first type of inputs providing absolute information to the CBAN, the second type provides ‘differential’ information, namely, the changes in the continuous variable. Sources of such inputs may be, for instance, self-generated movements providing velocity information to be integrated in the context of spatial navigation or sensory cues serving as pieces of evidence to be accumulated in the context of decision-making. In response to these inputs, the internal dynamics of the CBAN shifts the activity bump along the low-dimensional manifold—in a process akin to mathematical integration—such that the bump’s location reflects the value of the continuous variable. However, compared to the absolute information, the encoding accuracy of this integration process depends critically on an additional factor, namely, the integration gain of the network that relates the cumulative change in the continuous variable to the updating of the bump location in a proportional manner [27,28]. If this gain factor is miscalibrated, the result of the CBAN’s integration begins drifting away from the true value of the continuous variable; that is, it accumulates error. In the presence of absolute information sources such as visual landmarks for spatial localization, the aforementioned ‘basin’ mechanism continuously corrects error, preventing it from accumulating. However, without such absolute information, error accumulation continues, which may cause, for example, a CBAN integrating evidence to reach a decision threshold too soon or too late or a CBAN integrating an animal’s angular head velocity to over- or underestimate the correct head direction. Thus, a finely tuned integration gain is crucial for a CBAN to accurately encode a continuous variable based on inputs with only differential information.

Present CBAN models of path integration treat the integration gain as a constant that is perfectly set via carefully chosen, hard-wired model parameters (e.g., synaptic weights) [11, 29–31] (but see [32]). However, recent data from time cells and place cells of the rodent hippocampal formation, hypothesized to rely on CBANs [33], showed that the integration gain is actually a plastic variable whose value is adjusted based on the feedback from absolute information sources [34,35]. In the first study that demonstrated this phenomenon on place cells [35], the virtual visual landmarks, which provided the absolute information, were moved as a function of the animal’s movement. This movement created persistent error between the encoded location based on path integration and the actual location relative to the landmarks. Prolonged exposure to this conflict led to recalibration of the system’s integration gain, altering it in a direction and by an amount that reduced the positional encoding error. This recalibration was most evident after the landmarks were extinguished (i.e., when the absolute information to the putative CBAN was abolished); the space encoded by hippocampal cells during pure path integration either expanded or contracted, depending on the direction of the preceding landmark manipulation. Therefore, an open question is how CBANs adjust their integration gain based on error feedback from absolute information sources.

In the present paper, we aim to address this open question and generate testable physiological predictions about the neural mechanisms of gain recalibration in brain circuits that encode continuous variables. As a representative problem, we focus on hippocampal place coding and theoretically examine how visual landmarks, being the absolute information source, recalibrate the integration gain of a CBAN that encodes the animal’s position on a circular track, as demonstrated experimentally in [35]. To this end, we first derive analytical expressions of the integration gain, along with a simple dynamical model of the activity bump’s location in the ring attractor network, a specific type of CBAN for circular position encoding. In contrast to the previous work implicitly assuming the network’s integration gain as a constant, global parameter independent of the bump location in the low-dimensional manifold, our analysis reveals that the integrator gain of a CBAN is a spatially distributed, possibly inhomogeneous, parameter. We then employed control theory techniques to dissect the necessary algorithmic conditions for accurate recalibration of this spatially distributed integration gain via feedback from the absolute information sources. Mapping these conditions from the algorithmic level to the mechanistic level uncovered a key mechanistic requirement for gain recalibration. We found that, unlike correction of encoding errors that happens *automatically and implicitly* through network dynamics when feedback from an absolute information source is available, the process of learning/recalibrating the integration gain through Hebbian plasticity requires an additional neural signal that *explicitly encodes the error* in the network’s representation. In other words, all prior work demonstrating *error correction in a CBAN* does *not* require explicit error encoding, but our work shows that *correcting the integration process itself (i*.*e*., *recalibration)* requires an *explicit* representation of error. This error signal must be provided by changes in the firing rate of some neurons with one of two signals—the instantaneous error or the time-integral of the error—for recalibration of the integration gain. Finally, we propose a modified ring attractor network as an example CBAN model that instantiates our theoretical findings. Combining an error-rate code with Hebbian plasticity, this model achieves recalibration of integration gain in a CBAN, ensuring accurate representation for continuous variables.

## 2 Model Setup: Ring Attractor Network

A continuous bump attractor network is a recurrently connected neural network in which neighboring neurons excite one another and inhibit distant neurons according to a connectivity pattern known as local excitation and global inhibition [1, 38]. This connectivity gives rise to a persistent bump of activity as a stable, equilibrium state of the system. Invariance of the connectivity across the network leads to a continuum of such equilibrium states, called attractor states. Arrangement of neurons and the exact pattern of the recurrent connectivity determine the topology of this attractor. In the case of a ring attractor, neurons are arranged conceptually as a topological ring [11]. By sustaining an activity bump whose location can be shifted along the ring based on external inputs (i.e., relative and absolute information sources), a ring attractor network is well-suited to represent a variable on a closed curve (e.g., angular location of an animal on a circular track). Augmenting the arrangement of neurons to a two dimensional plane results in a plane attractor whose activity bump is well-suited to represent two variables, for example, the x and y coordinates of location in a room [37].

The activities of place and grid cells in 2D environments have been traditionally modeled using a plane attractor [12, 31, 37]. However, in the present investigation of gain recalibration based on location encoding, originally demonstrated in place cells from rats running laps on a 1D circular track, we chose the ring attractor as the basis of our model because of its analytical tractability.

The ring attractor that we analyzed is a network model consisting of three groups of neurons ordered in a ring arrangement: a central ring, a clockwise (CW) rotation ring, and a counter-clockwise (CCW) rotation ring [8, 11], as depicted in Fig. 1A. Neurons in this network receive synaptic input from other neurons in the network via intrinsic connections and from upstream neurons carrying velocity information (i.e., a ‘differential’ type of input) and the positional feedback from visual landmarks (i.e., an ‘absolute’ type of inputs) via extrinsic connections.

**Figure 1:**
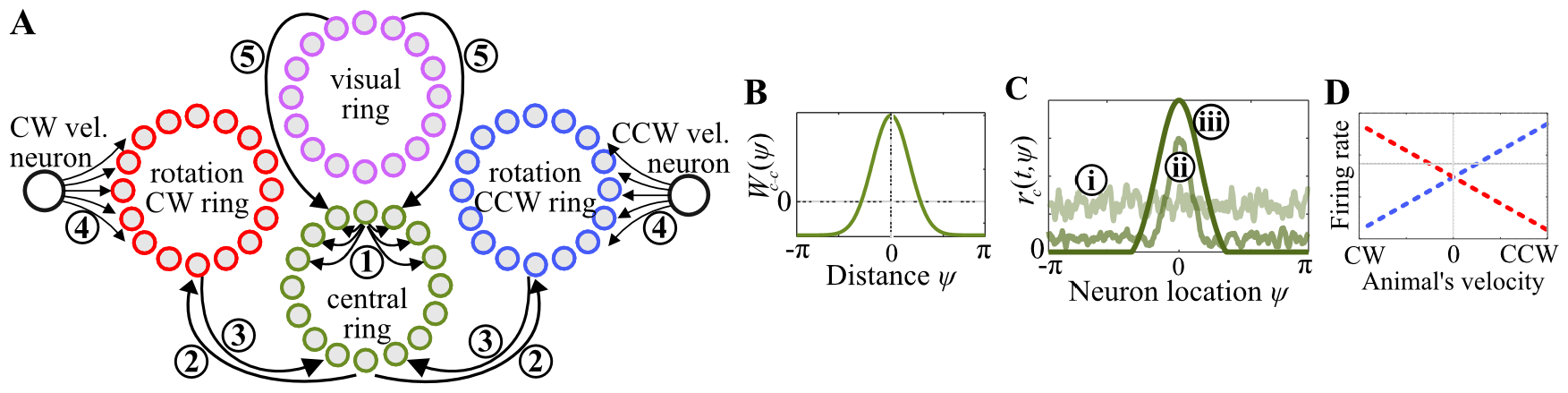
Ring attractor network model [10,31, 36, 37]. (A) Schematic representation of the model. The central ring forms the main body of the model based on its recurrent connections (labeled with ➀). Its reciprocal offset connections with the rotation CW and CCW rings (labeled with ➁ and ➂) creates a push-pull mechanism that modulates the intrinsically controlled neural activity of the central ring based on external inputs from the CW and CCW velocity neurons (labeled with ➃). An additional external input is provided to the central ring from the visual ring (labeled with ➄), corresponding to a set of sensory neurons that are tuned to visual landmarks. (B) Synaptic weight function *W*_c-c_ : *S*^1^ *→* ℝ that describes the recurrent connections within the central ring according to the well-known local excitation and global inhibition pattern. (C) Numerical demonstration of how recurrent connectivity within the central ring can autonomously maintain a persistent activity bump. Simulation of the central ring neurons was started with initial conditions that are assigned pseudo-randomly (light green line labeled with 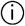 ). Within ∼100 milliseconds, a bump of activity emerges (medium green line labeled with 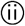 ). Eventually, the firing rates converge to an equilibrium, forming a persistent bump of activity (dark green line labeled with 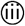 ). (D) Tuning curves of CCW and CW velocity neurons shown with blue-dashed and red-dashed lines, respectively.

We can model the dynamics of the ring attractor network using a set of equations that model the firing rate of a continuum of neurons in response to their synaptic inputs. If we parameterize a neuron based on its angle *ψ* ∈ *S*^1^ in the circular neural space, the model of the central ring neurons takes the form

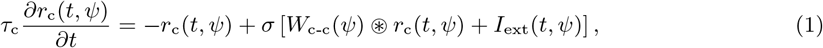

where *r*_c_(*t, ψ*) denotes the firing rate of the central ring neuron *ψ* at time *t, τ*_c_ denotes the synaptic time constant of central ring neurons, ⊛ denotes the circular convolution operation, *σ* denotes an activation function (chosen as rectified linear unit (RELU) in our current study), *I*_ext_(*t, ψ*) denotes external synaptic inputs to the central ring, and *W*_c-c_ : *S*^1^ *→* R denotes a rotationally invariant synaptic weight function that describes the recurrent connections (➀ in Fig. 1A) according to the pattern known as local excitation and global inhibition (Fig. 1B). Being a hallmark of the CBANs, this recurrent connectivity pattern leads to stabilization of a persistent “bump” of activity within the network [38–40] (Fig. 1C). While the shape of the emergent activity bump is determined by the shape of the recurrent connectivity pattern, the location of the bump can be controlled by external synaptic inputs to the central ring.

These external inputs are provided by neurons of the rotation rings and the visual ring. The rotation rings conjoin the self-movement velocity information with the positional information by receiving inputs from two different afferent neurons: By changing their firing rates in different directions (Fig. 1D), CW and CCW ‘velocity’ neurons signal the animal’s velocity information to their respective rotation rings through synaptic weight functions *W*_v-cw_, *W*_v-ccw_ : *S*^1^ *→* ℝ (➃ in Fig. 1A), whereas the central ring provides the positional information, represented by the bump location, to both rotation rings through synaptic weight functions *W*_c-cw_, *W*_c-ccw_ : *S*^1^ *→* ℝ (➁ in Fig. 1A). The resulting conjunctive codes of velocity and position within the rotation rings are then transmitted back to the central ring through *offset* synaptic weight functions *W*_cw-c_, *W*_ccw-c_ : *S*^1^ *→* ℝ (➂ in Fig. 1A), leading to a *shift* of the persistent activity bump within the central ring proportional to the animal’s velocity, a process known as path integration (PI). In contrast to the rotation rings, the visual ring does not explicitly receive any inputs; instead, its neurons are presumed to autonomously fire at specific locations of the animal relative to landmarks, capturing the absolute positional information received from visual landmarks available at each position (modeling how egocentric visual processes can calculate position from landmarks is beyond the scope of this paper). Through synaptic weight function *W*_vis-c_ : *S*^1^ *→* ℝ (➄, in Fig. 1A), this firing of the visual rings provides to the central ring a bump-like synaptic input encoding the animal’s “true” position relative to landmarks. This bump-like input pulls the activity bump of the central ring, hence correcting positional errors in the ring attractor’s representation.

Before concluding this section, we clarify an important distinction between traditional models of the ring attractor network and our model in the present paper. In traditional models, the synaptic weights *W*_v-cw_, *W*_v-ccw_ and *W*_cw-c_,*W*_ccw-c_ are treated as constants, each constrained to take a uniform value across the entire neural space. However, in our model, we intentionally relax this constraint. Instead, we treat these weights as functions that can vary throughout the neural space, possibly taking nonuniform values. While this approach may be less mathematically convenient, it becomes necessary when we later explore the possibility of gain recalibration through plasticity of spatially distributed synapses in the ring attractor network. As will be evident in the subsequent sections, our approach has broader implications, especially for the spatial metric of the ring attractor network. The complete mathematical model of the ring attractor, including the functional synaptic weights and the dynamics of the rotation and visual rings, is given in Appendix 6.1.

## 3 Algorithmic and Mechanistic Requirements for Gain Recalibration

In this section, we analyze the complex dynamics of our unconstrained ring attractor model to garner insight into how the network’s integration gain, hereafter referred to as the PI gain, can be recalibrated by visual landmarks. Our analysis begins with reduction of complex ring-attractor dynamics into a simple, one-dimensional differential equation model, including an analytical expression of the PI gain, in Section 3.1. Leveraging the analytical tractability of this simple model, we then identify algorithmic conditions for gain recalibration in Section 3.2. Finally, we use the analytical expression of the PI gain to map the algorithmic conditions within the simple model to mechanistic prerequisites for gain recalibration within the high-dimensional, complex model of the ring attractor network, in Section 3.3.

### 3.1 Dimensionality reduction reveals computational principles of the network

To derive a simple model for how the location of the attractor’s activity bump is controlled by external velocity and visual inputs, we follow the dimensionality reduction protocol described in [41]. Briefly, the protocol exploits the fact that ring-attractor dynamics constrain the population activity to form a bump whose overall shape stays invariant, but its center location can vary across the central ring. Although we do not know the exact solution to the network dynamics that can describe this activity pattern, we can ‘guess’ a solution form that describes its general properties without relying on a specific function. This guess, termed an *ansatz* solution, makes the analysis mathematically tractable by reducing the complex network dynamics to a one-dimensional differential equation that tracks the temporal change in the location of the activity bump as a function of external inputs.

Using this protocol, prior work derived the simple models for the traditional ring attractors, constraining the synaptic weights *W*_v-cw_, *W*_v-ccw_ and *W*_cw-c_,*W*_ccw-c_ to be constant [41, 42]. Here, we extend this approach to our unconstrained ring attractor. We develop the differential equation models progressively, starting from the simplest case that the animal is stationary in the absence of landmarks. Next, we add movement inputs and show that the network employs a spatially distributed PI gain. Finally, we extend the model to include landmark-based correction, and show that the network combines the spatially distributed integration with landmarks in a computation that resembles a Kalman filter [43]. The resulting model forms the basis for our search for the algorithmic conditions of the gain recalibration.

#### 3.1.1 Ansatz solution to network dynamics

When the central ring’s recurrent weight function *W*_c-c_ is symmetric with local excitation and global inhibition, firing rate of its neurons converges to a symmetric, persistent activity bump, taking nonzero values in a limited range and featuring a single peak corresponding to the internal representation of the animal’s position (Fig. 1C). Although specific functions such as thresholded Gaussians or sinusoids possess these characteristics and are often employed to explain neurophysiological data [11, 19, 44–47], we do not restrict ourselves to such a specific structure in the present analysis of the ring attractor. Instead, we assume that the firing rates *r*_c_(*t, ψ*) of the central ring neurons can be represented by a general ansatz solution

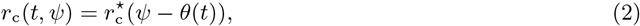

where 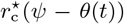 is a function that described the aforementioned persistent activity with a single peak at *ψ* = *θ*(*t*). This moderate generality allows our analysis to be valid for a broad range of ring attractor models with various particular ansatzes (including, but not limited to, the commonly used ones mentioned above). See Appendix 6.2.1 for a formal description of the properties of the ansatz function 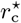.

The persistent activity in the central ring also spreads to the CW and CCW rotation rings through synaptic connections, resulting in the following ansatz solutions to their firing rates *r*_cw_, *r*_ccw_:

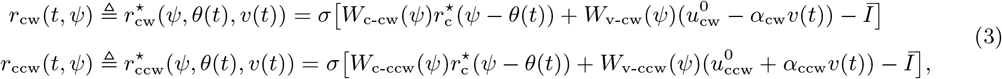

where 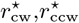 denote the assumed form of the ansatz solutions, 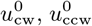 denote the baseline firing rates of the CW, CCW velocity neurons (during the animal’s immobility), *α*_cw_, *α*_ccw_ denote the absolute value of the slopes of the velocity neurons’ tuning curves (i.e., the absolute change in their firing rates per unit velocity of the animal), and *Ī* denotes global inhibition. With a properly set inhibition *Ī*, the solutions 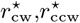 take the shape of a bump, similar to the ansatz 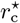 for the central ring (see Appendix 6.2 for further details).

Collectively, Eqs. (2) and (3) constitute a solution to the entire ring attractor network. That is, for a given velocity *v* of the animal, the firing rates of all neurons in a given network can be computed using the ansatz 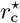 that describes the shape of the persistent activity bump within the central ring and *θ* that denotes the location of the bump. While the exact form of 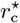 is determined by the profile of the recurrent weight function *W*_c-c_, the bump location *θ* is controlled by external inputs *I*_ext_ to the central ring. Therefore, assuming that the ansatz 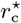 persists at all times like the previous work [41, 42], we can reduce the high dimensional ring-attractor dynamics to a one dimensional differential equation that models the position representation *θ* as a function of the external inputs. As derived in Appendix 6.2, if the central ring receives balanced (i.e., symmetric) inputs from the rotation rings during the animal’s immobility in the absence of landmarks (a classical assumption in CBAN models [3, 11, 31, 48, 49]), the position representation *θ* remains invariant. This invariance can be simply modeled as

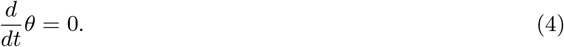

#### 3.1.2 Reduced-order path integration model reveals spatially distributed gain

In the present section, we examine how the position representation *θ*, decoded from the bump location, varies as the animal moves on a circular track in the absence of visual landmarks. By triggering differential changes in the firing of CW, CCW velocity neurons (Fig. 1D), such movement modulates the persistent activity bumps of the CW, CCW rotation rings also differentially. Synaptic connections from the rotation rings to the central ring then translate these changes in the activities of rotation rings to a change in the synaptic inputs to the central ring. As a result, the central ring no longer receives balanced excitation about its activity bump during the animal’s movement. With its magnitude proportional to the animal’s speed, this movement-based imbalance shifts the activity bump across the central ring, instantiating PI.

To obtain a model for the temporal dynamics of the position representation *θ* during this process, we apply the dimensionality reduction protocol described in [41]. Under mild assumptions detailed in Appendix 6.2.4, this reduction protocol leads to an ordinary differential equation

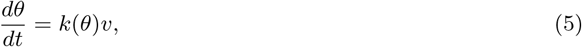

showing a linear relationship between the temporal change in *θ* and the animal’s velocity *v* via a factor *k*(*θ*). This factor quantifies the ring attractor network’s PI gain, and its analytical expression takes the form

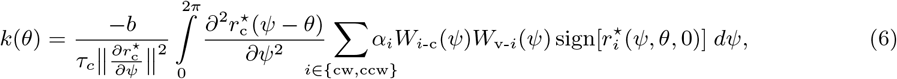

where *i* denotes the index of the summation, representing either the CW or CCW rotation ring, *α* denotes the absolute value of the slope of the velocity neurons’ tuning curves, and *b* denotes the value of the offset in the connections between rotation rings and the central ring. As shown by this equation, the PI gain *k*(*θ*) is a parameter determined by the network’s functional properties (i.e., profiles of the activity bumps and the tuning slope of the velocity neurons) and structural properties (i.e., time constant of the neurons and synaptic weights of the velocity-to-rotation ring and rotation-to-central ring connections), thereby capturing the complex interaction between the network’s internally generated persistent activity and the external inputs from the velocity neurons. A comprehensive examination of how these network properties relate to the PI gain is provided later in Section 3.3 when we explore the mechanisms of gain recalibration.

As mentioned previously, in our model, we do not adopt the assumption of the traditional models that constrains the synaptic weights of the velocity-to-rotation ring and rotation-to-central ring connections (*W*_v-cw_,*W*_v-ccw_ and *W*_cw-c_,*W*_ccw-c_, respectively) to be constants, each taking a uniform value in the entire neural space. Rather, we relax this constraint and treat these weights as functions that can vary along the neural space of the attractor network model, a heterogeneity likely to exist in biological networks. This functional treatment of the synaptic weights in our unconstrained model reveals an important characteristic of integration in CBANs: Unlike traditional treatments that implicitly assume a single, spatially global integration gain, CBANs employ a distributed, possibly inhomogenous, gain factor that can vary as the position representation varies (Equation 6 and top row in Fig. 2B). This implies for the ring attractor network that, by employing an inhomogenous PI gain, the network can adjust its spatial resolution locally, which would result in ‘overrepresentation’ or ‘underrepresentation’ of certain locations (bottom row in Fig. 2B) as is seen under various conditions of hippocampal place cell recordings [50, 51].

**Figure 2:**
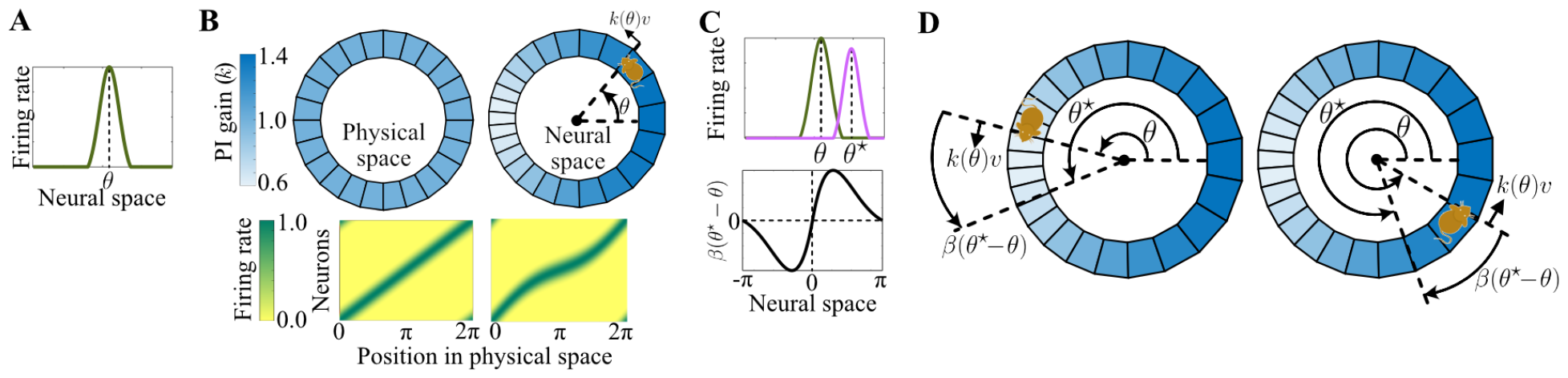
Models of the ring attractor’s position representation. (A) The position representation *θ*, decoded from the peak location of an example ansatz solution 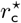 to the central ring’s firing rates. (B) Model of path integration. Top left: Circular track with uniformly spaced points. Top right: internal representation of this track with a spatially inhomogenous path-integration (PI) gain that ranges from 0.6 at *θ* = *π* to 1.4 at *θ* = 0. The position representation *θ* is visualized here by the rat. As the rat moves through physical space at velocity *v*, the representation moves through neural space at *k*(*θ*)*v*. Bottom: Firing rate of uniformly distributed cells in the neural space as a function of the animal’s position in physical space. Left shows a ‘traditional’ network model, including a single, global PI gain of 1. Right shows our unconstrained network model with the spatially inhomogenous PI gain in the top row. (C) Stabilizing visual feedback. Top: The central ring’s activity bump 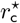 (green) and the bump-shaped synaptic input *I*_vis_ to the central ring from the visual ring (pink). The activities of both rings are aligned with respect to the same neural space, so in this example, the visual ring bump is “ahead of” the central ring bump. Bottom: The function *β* captures the stabilizing feedback from visual landmarks. Note that *β* operates on *the difference, θ*^⋆^ *− θ*, so here, the *x* axis is a dummy variable. (D) Model of path integration with visual feedback. Left: the temporal change in the position representation is visualized as a cartoon rat, symbolizing the current representation as in (B), pulled by two arrows, one corresponding to updating by the path integration term *k*(*θ*)*v* and the other corresponding to updating by the visual feedback term *β*(*θ*^⋆^ *− θ*). Note that in this position, the PI gain is “low” and thus PI underestimates position relative to the landmarks. Right: Same as Left but PI overestimates position relative to the landmarks due to “high” PI gain in this position.

#### 3.1.3 Landmark correction to path integration resembles a Kalman filter

Lastly, we examine how the position representation *θ* varies during the most general case that the animal moves on the circular track in the presence of landmarks. To derive this one-dimensional model of *θ*, we employ the same dimensionality reduction protocol [41] but this time taking into account the visual neurons that provide feedback from landmarks. The resulting model will be crucial later when searching for the algorithmic conditions of the gain recalibration.

It is known that feedback from landmarks anchors the internal representation of position in a stable manner [22–25]. In classical ring attractor models, this stabilizing feedback is achieved with an allocentrically anchored, bump-like synaptic current applied onto the central ring from the visual ring, encoding the animal’s location relative to the landmarks [52,53]. We adopt the same approach in our unconstrained ring attractor model and incorporate the following synaptic current from the visual ring onto the central ring:

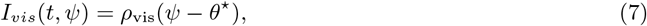

where *ρ*_vis_ : *S*^1^ *→* ℝ denotes a function describing the bump shape of this current, and *θ*^⋆^ denotes the location of the peak current, corresponding to the animal’s current position relative to landmarks. Note that, throughout the paper, we use the superscript ^⋆^ to distinguish a true value of a quantity (measured relative to the external world) from its internal value (e.g., *θ*^⋆^ vs *θ*).

With Equation (7) in mind, applying the dimensionality reduction protocol [41] leads to a differential equation model for how the position representation *θ* is controlled by the PI and the landmarks as follows:

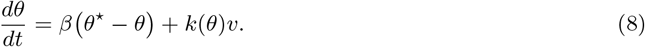

Here, *β* : *S*^1^ *→* ℝ^1^ is a function that, locally, takes the same sign as its argument (Fig. 2C). This sign property, a form of negative feedback, is crucial for stable control of the position representation *θ*; for example, in the absence of movement, the sign property of *β* ensures *θ → θ*^⋆^. In the more general case, when the animal’s velocity is nonzero, the position representation *θ* is changed by a combination of the (stabilizing) visual feedback *β* : *S*^1^ *→* ℝ and the feedforward PI-related term *k*(*θ*)*v*, as given in Equation (8). While the PI-related term, *k*(*θ*)*v*, shifts the position representation proportionally with the animal’s velocity, the function *β* acts in an additive manner to bring *θ* toward *θ*^⋆^, the animal’s true position relative to visual landmarks. This fusion between the feedforward PI-related term and feedback correction from landmarks is illustrated in Fig. 2D. A properly balanced combination would stabilize *θ* around *θ*^⋆^, with only minor deviations due to PI error. As a result, the ring attractor effectively “estimates” the position of the animal, a computation that bares striking similarity to a Kalman filter in engineering. Here, the state being estimated by the network is the animal’s true position, *θ*^⋆^, and the estimate is the bump location, *θ*.

### 3.2 Control Theory Reveals Algorithmic Conditions for Gain Recalibration

Experiments showed that the PI gain is a plastic variable that can be recalibrated accurately by visual landmarks [35]. In this section, we seek necessary conditions for this recalibration using Equation (8), the simple model presented in the previous section.

We begin by recalling the experimental conditions that brought about the recalibration of the PI gain [35]: An animal moved on a circular track while an array of visual landmarks was rotated around the track as a function of the animal’s velocity and an experimentally controlled, visual gain factor, *k*^⋆^. When *k*^⋆^ *<* 1, the landmarks moved in the same direction as the animal, decreasing the animal’s speed relative to the landmarks; when *k*^⋆^ *>* 1, the landmarks moved in the opposite direction as the animal, increasing the animal’s speed relative to the landmarks; when *k*^⋆^ = 1 (veridical condition), the landmarks were stationary. Neural recordings showed that persistent exposure to these visual conditions recalibrated the animal’s PI gain such that a tight correlation was observed between the average value of the PI gain measured over many laps after the landmarks were extinguished and the final value of the visual gain *k*^⋆^ before the landmarks were extinguished. Here, we analyze this recalibration using our simplified ring attractor model.

The experimental conditions leading to the gain recalibration can be simulated in a ring attractor model with a visual synaptic drive *I*_vis_ revolving around the central ring at a rate equal to the animal’s velocity *v* times the visual gain *k*^⋆^. Extending (8), the simple model for the position representation *θ*, with an additional equation for modeling the changes in the peak location *θ*^⋆^ of such a visual drive, we obtain

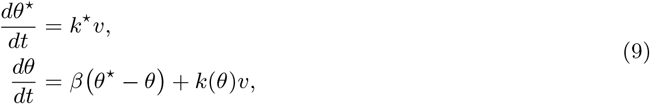

where the first equation models the relation between the visual gain *k*^⋆^, the animal’s velocity *v*, and the resulting temporal change in the visual drive’s bump location *θ*^⋆^, and the second equation models the change in the attractor’s bump location *θ* based on the external velocity and visual inputs. Recall that, as we previously showed, the PI gain *k* of a ring attractor network is a spatially distributed parameter that may potentially take different values as the network’s position representation *θ* varies during the animal’s movement in the environment. However, in recalibration experiments [35], average neural activity over many laps was used to estimate the average value of the PI gain, providing no information about whether the PI gain took different values across the environment as predicted by the model. Therefore, from a theoretical perspective, the experimental result that the PI gain was recalibrated to the visual gain [35] only suggests that its *spatial average k*_0_ converges to the visual gain *k*^⋆^ (i.e., lim_t*→∞*_ *k*_0_(*t*) = *k*^⋆^), where

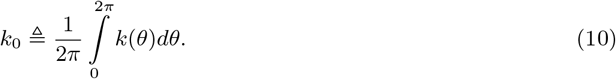

This convergence of *k*_0_ toward *k*^⋆^ necessitates *k*_0_ to be updated over time during recalibration. The firing rate of neurons in the network is a biologically plausible candidate for controlling these updates. Therefore, to garner mathematical insight into this process, we searched for a general equation that could represent the updating of *k*_0_ based on firing rates within the ring attractor, assuming an environment with a spatially homogenous feedback from visual landmarks. Under mild assumptions described in Appendix 6.3.1, this search led to a surprisingly simple equation

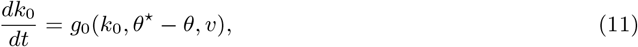

where *g*_0_ : ℝ *× S*^1^ *×*ℝ *→* ℝ denotes an analytic function that instantiates the instantaneous change in *k*_0_ based on three variables: the current gain *k*_0_, the animal’s velocity *v* and the difference between the visual drive’s position representation *θ*^⋆^ and the ring attractor’s position representation *θ, without* directly depending on either *θ* or *θ*^⋆^ (since the visual feedback is assumed to be uniform across the environment).

#### 3.2.1 Necessity of matching the sign in gain change with the product of error and velocity

The update rule *g*_0_ could be any function fitting the form in Equation (11); however, some such functions may fail to result in gain recalibration, i.e., the PI gain’s spatial average *k*_0_ would not converge to the visual gain *k*^⋆^. What are the necessary properties of the gain update rule *g*_0_ for convergence of *k*_0_ to *k*^⋆^?

To seek these properties, we revisit Equation (9), the simple model of the central ring’s and visual ring’s position representations *θ* and *θ*^⋆^, now taking into account that the PI gain’s spatial average *k*_0_ is time varying according to the gain update rule in Equation (11). Perfect convergence of *k*_0_ to *k*^⋆^ through this update rule would imply that the error between these two gains, namely 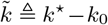, approaches zero. When this gain error becomes zero, it is intuitively expected that the error in the attractor’s position representation relative to the visual drive, namely 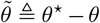, also approaches zero. Hence, analyzing the temporal progression of these error terms— namely the gain error 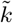 and the positional error 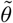 —provides an opportunity to garner insight into the algorithmic underpinnings of the gain recalibration process.

To this end, let us assume a constant visual gain, implying *dk*^⋆^*/dt* = 0. To track how the PI gain’s spatial average *k*_0_ recalibrates to this constant value, we subtract the second row of Equation (9) from its first row and Equation (11) from *dk*^⋆^*/dt* = 0. This yields the so-called *error dynamics*

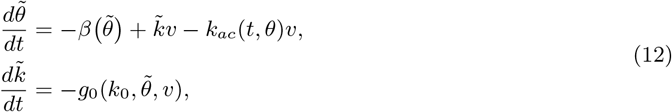

where *k*_ac_(*θ*) ≜ *k*(*θ*) *− k*_0_ captures the spatial variation of PI gain *k*(*θ*) from its spatial average *k*_0_. Analyzing these error dynamics with tools from feedback control theory, we then identify the necessary conditions for complete recalibration of *k*_0_ to *k*^⋆^.

First, we find that, if the ring attractor achieves and maintains zero gain error 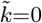, as required for complete recalibration, then it also maintains zero positional error 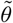, rendering the origin 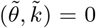 an equilibrium point of the error dynamics. Convergence to this equilibrium point, howewer, requires the animal must be moving, else there exists no gain update rule that can achieve it. This is an unsurprising result, because, were the animal stationary, the visual landmarks would correct all the positional error 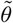, making the gain error 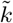 imperceptible to the animal. Assuming accordingly that the animal is always moving, our analysis then reveals a sign requirement that must be satisfied by any gain update rule *g*_0_. For stable convergence of the gain and positional error to the equilibrium point at the origin 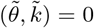, the gain’s spatial average *k*_0_ must be updated in the same direction as the product of the animal’s velocity *v* and the ring attractor’s positional error 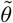 in some neighborhood of 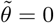:

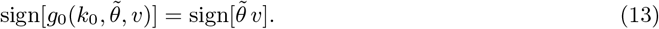

See Appendix 6.3.2 for formal statements and proofs of these findings.

What if the biological system can be recalibrated only partially? That is, at the steady state, the PI gain’s spatial average *k*_0_ converges to a value biased towards, but not necessarily the same as, the visual gain *k*^⋆^. Under a simplifying assumption that the animal’s velocity is constant, we can generalize our findings for the complete recalibration to a case that covers both complete and partial recalibration. To this end, we re-analyze the error dynamics (Equation 12). First, we find that convergence of the gain error 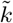 to some value, say 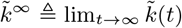, after recalibration results in convergence of the ring attractor’s positional error 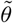 also, say to 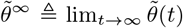. If the gain recalibration is complete (i.e., 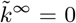), then this steady-state positional error 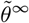 is zero; if the recalibration is partial (i.e., 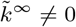), however, 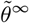 is nonzero, proportional to the product of the steady-state gain error 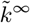 and the animal’s velocity *v*. Our analysis then reveals a generalized sign requirement that must be satisfied by any gain update rule: For gain recalibration, the gain’s spatial average *k*_0_ must be updated in the same direction as the product of the animal’s velocity *v* and the deviation of the current positional representation error 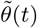 relative to its steady-state value 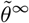 in some neighborhood of 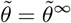:

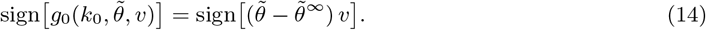

This generalized sign requirement captures both complete and partial recalibration; it, for example, reduces to Equation (13), the requirement for the complete gain recalibration, if the steady-state positional error 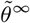 is zero. See Appendix 6.3.3 for formal statements and proofs of these findings.

#### 3.2.2 Sufficiency of positive gain change with respect to the product of error and velocity

We next analyze whether a gain update rule satisfying Equations (13) and (14), the necessary algorithmic conditions for recalibration, achieves gain recalibration, i.e., is it sufficient?

To address this question, we further analyzed the error dynamics in Equation (12). This analysis reveals a sufficient condition for gain recalibration. According to this condition, a gain update rule is guaranteed to achieve gain recalibration (may be partial or complete) for any visual gain *k*^⋆^ and a given velocity *v* of the animal if it has a positive slope with respect to the product of velocity *v* and positional error 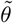 at its zero value (i.e., 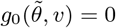), namely,

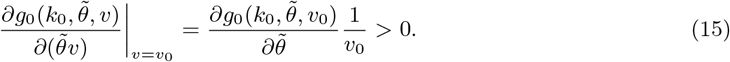

This sufficient condition is a small extension to the generalized necessary condition for recalibration such that it guarantees recalibration if a gain update rule satisfies the generalized sign condition in Equation (14) together with the mild additional requirement that the update rule also has a nonzero derivative with respect to the product of 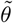 and *v*. For example, a linear update rule 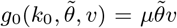 with *μ >* 0 satisfies the sufficient condition but a cubic rule 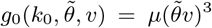 only satisfies the necessary condition. We now would like to demonstrate how this sufficiency leads to gain recalibration via two example update rules. Although both rules satisfy the sufficient condition, Example 1 achieves complete recalibration, while example 2 achieves only partial recalibration.

##### Example 1.

*The simplest gain update rule that satisfies Equation (15), the slope condition guarantee for gain recalibration, takes the form*

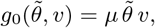

*where μ denotes a positive learning rate. Furthermore, the update rule takes the same sign as the product* 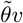, *thus also satisfying the necessary condition for complete recalibration (Fig. 3B)*.

**Figure 3:**
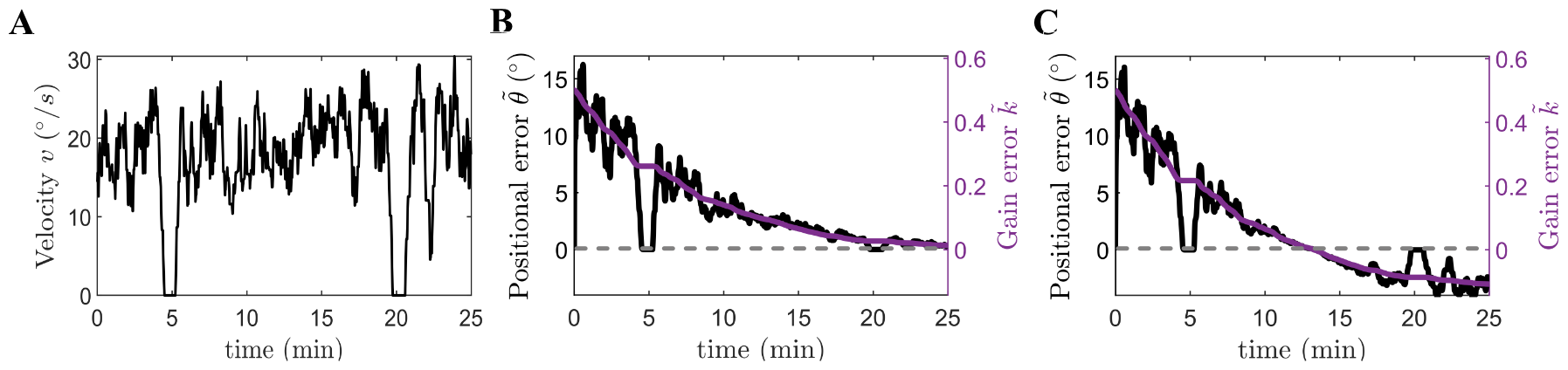
Numerical simulation of the two example gain update rules. For both simulations, we chose the initial condition *k*_0_(0) = 1 and the parameters 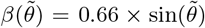, *k*^⋆^ = 1.5, *μ* = 0.02. The gain choices imply that the initial value of gain error is 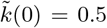. Additionally, we chose *η* = 0.12 for the second example. (A) Temporal progression of the smoothed animal’s velocity from an experiment in [35]. (B) Simulated error trajectories under example gain update rule 1. As soon as the animal begins its movement at *t* = 0, the positional error 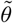 (black line relative to the left y axis) quickly increases because of the nonzero gain error 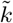 (purple line relative to the right y axis). As the animal recalibrates its gain, the gain error gradually converges to zero (i.e., 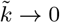), accompanied by positional error gradually converging to zero also (i.e., 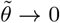). In addition to these gradual convergent trends, the error trajectories include many fast, transitory changes. As can be seen from the black line, the instantaneous value of positional error 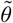 is correlated with the animal’s velocity *v* also. For example, when animal slows down, the positional error decreases, becoming zero when the velocity is zero. This is a reflection of the relatively increased landmark stabilization 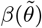 when path integration inputs *kv* are decreased. On the other hand, the temporal changes in the gain are correlated with the multiplication of the positional error 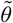 and the animal’s velocity *v* as determined by the gain update rule *g*_0_. When the animal pauses temporarily around minutes 5 and 20 (i.e., *v* = 0), the positional error 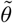 is completely corrected by landmarks (i.e., 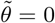), causing the gain updates to pause also (i.e., 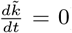). As the animal continues moving, the positional error and the velocity fine-tune the gain until the gain error converges to zero, demonstrating that the system can achieve complete gain recalibration. (C) Simulated error trajectories under example gain update rule 2. The convention is the same as panel B. The error trajectories exhibit similar trends to the panel B except that their final values do not converge to zero, demonstrating that the system can only achieve partial gain recalibration.

##### Example 2.

*This example is inspired from the modified ring attractor network that we propose in Section 4 as a model that achieves gain recalibration. Let μ denote a positive learning rate as before and η denote a constant. Consider the gain update rule*

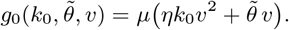

*Since its partial derivative is positive (i*.*e*., 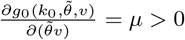*), this update rule results in gain recalibration like Example 1. However, unlike Example 1, the recalibration can be complete or partial, depending on the value of η. If η* = 0, *the update rule takes the same sign as the product* 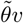, *thus satisfying the necessary condition for complete recalibration. Otherwise, it only satisfies the necessary condition for partial recalibration by taking the same sign as* 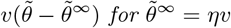 *(Fig. 3C)*.

### 3.3 Mechanistic Constraints Reveal Instrumental Role of Positional Error Codes

What are the mechanistic prerequisites for meeting the necessary algorithmic condition derived in the previous section for gain recalibration? To adddress this question, we investigate an analytical expression of the PI gain’s spatial average *k*_0_ which can be simply obtained by averaging the PI gain *k*(*θ*) in Equation (6) over *θ* as follows:

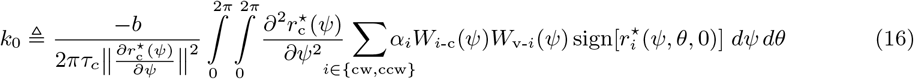

The terms in this expression identifies a number of possible loci or mechanisms for updating *k*_0_ that satisfy the algorithmic requirement for gain recalibration in Equation (13). These terms include: (i) the offset *b* in the central-to-rotation ring connections, (ii) the synaptic time constant *τ*_c_, (iii) the slope parameters *α*_cw_,*α*_ccw_ quantifying the absolute value of the tuning slopes of velocity neurons, (iv) the synaptic weight functions *W*_v-cw_,*W*_v-ccw_ of the velocity-to-rotation ring connections, (v) the synaptic weight functions *W*_cw-c_,*W*_ccw-c_ of the rotation-to-central ring connections, (vi) the function 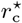 describing the central ring’s persistent activity bump, and (vii) the functions 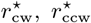 describing solutions to the rotation ring’s persistent activity bump. Note that we treat the CW and CCW components of the same term as an inseparable pair. Out of the seven terms, we consider the last five terms (iii-vii) as candidates driving the gain recalibration within the ring attractor model via temporal changes, implicitly assuming that the first two terms, the offset *b* and the synaptic time constant *τ*_c_, are “hardwired” (i.e., time-invariant).

The rationale behind excluding the first two terms arises, in part, from limitations of our modeling framework. First, the rate-based approach, upon which we describe the network dynamics, does not include any cellular and receptor details to capture possible temporal changes in the synaptic time constant *τ*_c_. Instead, our model includes *τ*_c_ as a “lumped parameter” reduction of complex phenomena that governs the changes in membrane potential with ion flux through receptors; future work could use chemical kinetics modeling to investigate how changes in *τ*_c_ could contribute to gain recalibration, but that is beyond the scope of the present study. Second, our model employs a simplified one-to-one connectivity between the rotation rings and the central ring such that one neuron in a rotation ring connects to only one neuron in the central ring with a fixed offset *b*, rather than a one-to-all connectivity required to capture plasticity in *b* through gradual modulation of weights along the neural space. Therefore, excluding *τ*_c_ and *b* from further consideration, we focused our analysis on the remaining five terms as the driver of gain recalibration.

By analyzing the relation between the temporal change in each of these candidate terms and the resulting temporal change in *k*_0_, we find that rate-based encoding of the positional error 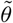 is required to satisfy the necessary algorithmic condition for the gain recalibration regardless of which term drives the changes in PI gain. However, the specific nature of the error code depends on the driver term (Fig. 4A) as demonstrated in the next subsections.

**Figure 4:**
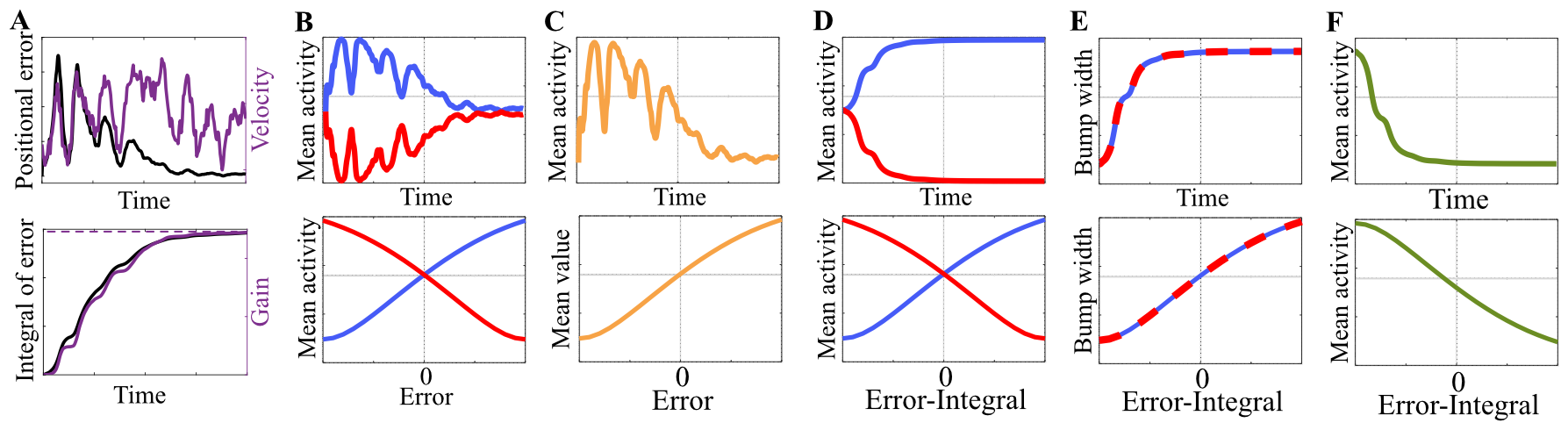
Visualization of mechanistic constraints for a numerical simulation based on a hypothetical gain update rule 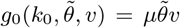 Except for the animal’s velocity profile, we chose the parameters and initial conditions the same as Fig. 3B. For color coding, we use Fig. 1A as the reference, where red and blue denote the CW and CCW rotation rings, and green denotes the central ring. (A) Top graph shows the simulated velocity of the animal (purple line) on the right y axis and the temporal progression of the positional error (black line) on the left y axis. Notice the synchronous fluctuations in the positional error and the animal’s velocity. As explained in Fig. 3B, these synchronous fluctuations occur because the positional error is correlated with the animal’s velocity. Bottom graph shows the PI and visual gains with solid and dashed purple lines, respectively, on the right y axis and the time-integral of the positional error with the black line on the left y axis. Notice that as the PI gain gradually converges to the visual gain, the temporal progression of the time-integral of the positional error follows a very similar trajectory. This similarity indicates that the integration gain reflects the past accumulation of positional representation errors, thus opening up the possibility for the network to track the time-integral of the error as a proxy signal to encode the integration gain. (B) The mechanistic constraint for recalibration through plasticity of the velocity-to-rotation ring connections. Top graph shows the mean firing rates of the CCW and CW rotation rings over time with blue and red lines. Notice that they are similar to the trajectory of the positional error in panel A, except that the changes in the CW rotation ring’s mean firing rate (red line) is the negative of those in the CCW rotation ring’s mean firing rate. Bottom graph shows the direct relationship between mean firing rates and positional error in the attractor’s representation. (C) The mechanistic constraint for recalibration through plasticity of the rotation-to-central ring connections. Top graph shows the mean firing rates of either the rotation rings or the central ring over time with the orange line. Notice that the changes in these firing rates follow a similar trend as the temporal progression of the positional error. Bottom graph shows this relationship directly (the positive correlation is chosen arbitrarily as our analysis does not provide a conclusive insight into the required direction). (D) The mechanistic constraint for recalibration through changes in the velocity neurons’ slopes. Top graph shows the mean firing rate of the CCW and CW rotation rings, the same quantities as panel B. However, unlike panel B where the mean firing rates were similar to the instantaneous positional error, the mean firing rates in this panel are similar to the time-integral of the error. Bottom graph shows this relationship between the mean firing rates and the time-integral of the error directly. (E) The mechanistic constraint for recalibration through changes in the rotation rings’ activity bumps. Top graph shows the bump width of both rotation rings over time. Similar to how the mean firing rates of the rotation rings encode the time-integral of the positional error in panel D, the bump widths encode the time-integral of the error in this panel. Bottom graph shows this relationship directly. (F) The mechanistic constraint for recalibration through changes in the central ring’s activity bump. Top graph shows the temporal progression of the mean firing rate of the central ring, which is tightly but negatively correlated with the temporal progression of the time-integral of the positional error. Bottom graph shows this relationship directly.

#### 3.3.1 Plasticity in the velocity pathway requires a rate code of the instantaneous error

We first consider the scenario that the temporal change in the PI gain is driven by temporal changes in either set of the synaptic weights along the pathway from velocity neurons to the central ring. These sets include the pair *W*_v-cw_, *W*_v-ccw_, describing the strength of velocity-to-rotation ring connections (➃ in Fig. 1A), and the pair *W*_cw-c_, *W*_ccw-c_, describing the strength of rotation-to-central ring connections (➂ in Fig. 1A).

According to Equation (16), the CW and CCW components of these weight pairs additively influence the PI gain *k*_0_. This additive influence suggests a possibility where CW and CCW components vary independently of one another during recalibration to tune the PI gain’s spatial average *k*_0_. However, this possibility is limited by the requirement that the central ring must receive balanced (i.e., symmetric) inputs from the CW and CCW rotation rings to keep the activity bump stationary during the animal’s immobility as we show in Appendix 6.4. Thus, we assume hereafter that the CW and CCW components vary in a coordinated fashion to ensure that their individual contribution to *k*_0_ is symmetric. This symmetry assumption implies that if the overall value of *k*_0_ changes as per the necessary algorithmic condition in (13), then the individual contribution to this change from both CW and CCW components must be in the direction of the product of the animal’s velocity *v* and the positional error 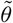. To identify the mechanistic underpinnings of such symmetric gain recalibration, we revisit Equation (16). By differentiating this equation with respect to time and considering Hebbian plasticity as the mechanism underlying the changes in the weight pairs *W*_v-cw_, *W*_v-ccw_ or *W*_cw-c_, *W*_ccw-c_, we find that the algorithmic condition translates into a mechanistic constraint as follows:

##### Hebbian plasticity of the velocity-to-rotation ring connections (W_v-cw_, W_v-ccw_)

A change in the weights *W*_v-cw_, *W*_v-ccw_ leads to a commensurate change in the speed at which the network’s activity bump is shifted along the ring for a given speed of the animal. These commensurate changes suggest a positively correlated relationship between the weights *W*_v-cw_, *W*_v-ccw_ and the PI gain *k*_0_. Assuming the symmetry between CW and CCW components, we indeed show in Appendix 6.4.1 that Equation (16), relating the weights *W*_v-cw_, *W*_v-ccw_ to the PI gain *k*_0_ via positively-weighted integrals, can be reformulated as

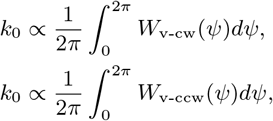

where the *∝* symbol denotes the existence of a positively-sloped, proportional relationship. Because of these positive correlations, satisfying the algorithmic condition for recalibration implies that the average strength of both CW and CCW velocity-to-rotation ring synapses is modified in the direction of the product of the animal’s velocity (*v*) and the network’s positional error 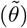, namely,

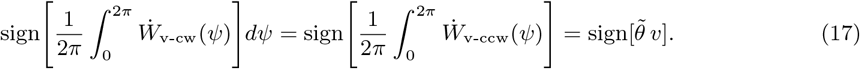

Here, the dots appearing over the weights denote the temporal change in the weight. Recall that Hebbian plasticity of a synapse is driven by the joint activity of pre- and post-synaptic neurons. In the specific case of velocity-to-rotation ring synapses, the pre-synaptic side is composed of the velocity neurons, designated to solely encode the animal’s velocity *v* with tuning curves that have a negative slope for the CW velocity neuron and a positive slope for the CCW velocity neuron (previously shown in Fig. 1D). Therefore, in a manner matching these differential signs of *v*-encoding on the presynaptic side, the CW and CCW rotation rings on the post-synaptic side *must monotonically decrease and increase their average firing rates with the instantaneous positional error* 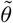 to satisfy the equality in Equation (17) (Fig. 4B). Mathematical details are provided in Appendix 6.4.1.

##### Hebbian plasticity of the rotation-to-central ring connections (W_cw-c_,W_ccw-c_)

Like the velocity-to-rotation ring connections discussed above, the rotation-to-central ring synaptic weight functions *W*_cw-c_,*W*_ccw-c_ enter linearly in the calculation of the PI gain in Equation (16). Therefore, like the previous case, the algorithmic condition for recalibration via *W*_cw-c_,*W*_ccw-c_ requires Hebbian plasticity to modify their average strength in the direction of the product of the animal’s velocity (*v*) and the network’s positional error 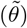, namely,

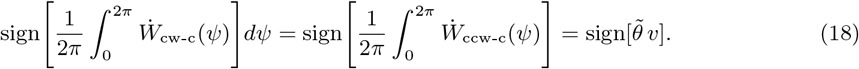

In the previous case, the mechanistic prerequisite for meeting a similar sign requirement was error-encoding on the *postsynaptic* side since the pre-synaptic neurons were assumed to solely encode the velocity. In the present case, however, it is feasible to encode the error in either the *pre-* or *post-synaptic* side since neither side is subject to such a limitation. Therefore, when the animal is traveling in one direction (as was the case in the experiments that originally demonstrated the gain recalibration [35]), satisfying the equality in Equation (18) requires mean firing rate of either the rotation rings or the central ring *vary monotonically with the network’s instantaneous positional error* (Fig. 4C). However, unlike the previous case, our analysis of the present case does not provide conclusive information about the direction of these monotonic relations. Mathematical details are provided in Appendix 6.4.2.

Collectively, these findings show that Hebbian plasticity in the pathway carrying the external velocity information to the central ring requires a rate code of the network’s instantaneous positional error to update the synaptic weights in the direction of the product of the animal’s velocity and the error as per the algorithmic condition for gain recalibration in Equation (13).

#### 3.3.2 Plasticity elsewhere requires a rate code of the time-integral of the error

We next consider the scenario that the synaptic weights along the pathway from velocity neurons to the central ring are hardwired (i.e., constant). This scenario implies that the gain recalibration is instead driven by temporal changes in one of the three *firing-rate* related terms, including the slope parameters *α*_cw_, *α*_ccw_ quantifying the absolute value of the slopes of the CW, CCW velocity neurons’ tuning curves, the ansatz functions describing the persistent activity bumps 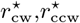 of the rotation rings, or the ansatz of the persistent activity bump 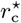 of the central ring. Independent of which of these terms undergoes temporal changes, the algorithmic condition for gain recalibration translates into a mechanistic constraint that, while still a rate code of error, differs in its fundamental characteristics as detailed below:

##### Changes in the slopes of velocity neurons’ tuning curves (α_cw_, α_ccw_)

As shown previously in Fig. 1D, the CW and CCW velocity neurons are tuned to the animal’s velocity with slopes having different signs. Assuming these differential signs to be hard-wired (i.e., constant), we examine in the present case that absolute values of the slopes, denoted by parameters *α*_cw_ and *α*_ccw_, undergo temporal changes. A change in these parameters leads to a commensurate change in the speed at which the network’s activity bump is shifted along the central ring for a given speed of the animal. As in the previous section, this implies a positively correlated relationship between the slope parameters *α*_cw_ and *α*_ccw_ and the PI gain *k*_0_. Indeed, this relationship is explicitly seen in Equation (16). Thus, as in the previous section, satisfying the algorithmic condition for recalibration (Equation (13)) requires *α*_cw_ and *α*_ccw_ to *change* in the direction of the product of the animal’s velocity (*v*) and the network’s positional error 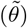, namely,

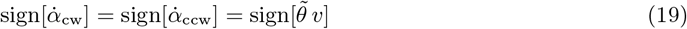

An implication of this requirement is that, when the animal is traveling in one direction, the *change in the slope parameters* is monotonically related to the positional error, reflecting its value on a moment-to-moment basis with a sign additionally depending on the sign of the velocity. When these changes are integrated over time, *the current value of the slope parameters* reflects the accumulation of positional errors through the past, depending monotonically on the time-integral of the positional error in a direction that depends on the animal’s velocity. Through connections between the velocity neurons and the rotation rings, these monotonic relationships are also translated to the mean firing rate of rotation rings. The direction of the monotonic relationships between the time-integral of the error and the mean firing rate of the rotation rings, however, depends additionally on the sign of the velocity neurons’ tuning slopes. For example, when the animal is traveling in one direction (say CCW) like the original recalibration experiments [35], the mean firing rate of the CCW rotation ring increases monotonically with the time-integral of the error due to the positive slope of the CCW velocity neuron. Conversely, the mean firing rate of the CW rotation ring decreases monotonically with the time-integral of the error due to the negative slope of the CW velocity neuron (Fig. 4D). If the animal travels in the other direction, the direction of these monotonic relationships is also reversed. Mathematical details are provided in Appendix 6.4.3.

##### Changes in the persistent activity bump of the rotation rings 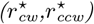

Velocity information is transmitted to the central ring by the rotation rings whose firing rates are modulated by the animal’s velocity. In this transmission, an increase in the number of actively firing rotation ring neurons (i.e., larger width of the bumps 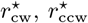) would result in a commensurate increase in the number of central ring neurons that receive the velocity information. As a result, the network’s activity bump shifts faster along the central ring even when the animal’s velocity is unchanged. This implies a positively correlated relationship between the widths of the rotation rings’ activity bumps and the PI gain *k*_0_, reminiscent of the relationship in the previously investigated case of *α*_cw_,*α*_ccw_. Thus, like the previous case, satisfying the algorithmic condition requires *the rotation rings’ activity widths to change monotonically* with the product of the animal’s velocity and the positional error. Consequently, when the animal is traveling in one direction (say positive), the widths of the rotation rings’ activity bumps must *increase monotonically with the time-integral of the error* (Fig. 4E). If the animal’s travel direction is negative, this relationship turns into a monotonically decreasing function. Mathematical details are provided in Appendix 6.4.4.

##### Changes in the persistent activity bump of the central ring 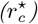

Consider as an example that there are two networks with the same Gaussian bump profile but one has a higher peak firing rate. In this case, if all else is equal, the network with the higher firing requires higher velocity inputs to shift its activity bump from point A to point B in the same time as the other network. This need of higher velocity inputs indicates an inversely correlated relationship between the magnitude of the central ring’s activity bump and the PI gain *k*_0_, which can also be verified from Equation (16) wherein the denominator includes a term proportional to the bump magnitude: the squared norm of the activity bump’s gradient. Thus, satisfying the algorithmic condition for gain recalibration is subject to a mechanistic constraint that is similar to the previous case in spirit but slightly different due to the inverse effect: When the animal is traveling in one direction (say positive), the mean firing rate of the central ring must *decrease monotonically with the time-integral of the error* (Fig. 4F). If the animal’s travel direction is negative, the direction of this monotonic relationship is reversed. Note that, for deriving this result, we assume the general shape of the central ring’s activity bump to be invariant, unlike its magnitude. Mathematical details are provided in Appendix 6.4.5.

Collectively, these findings show that gain recalibration might be possible without synaptic plasticity in the pathway carrying the external velocity information to the central ring, provided that there is a rate code of the time-integral of the positional error as opposed to the error itself. Our analysis does not provide any insights into the mechanisms necessary for such a rate code and the temporal changes in the terms associated with it (e.g., slopes of velocity neurons). However, it may be the case that plasticity elsewhere than the velocity pathway is required. Independent of the underlying mechanism, however, an absence of plasticity in the velocity pathway would lead to the conclusion that the PI gain is no longer encoded in the synaptic weights: instead, it is encoded in the firing rates that track the time-integral of the error, a proxy of the PI gain (bottom row in Fig. 4A).

## 4 Implementing Gain Recalibration in a Ring Attractor

In this section, we propose a modified ring attractor model that can achieve gain recalibration through a mechanism devised based on the theoretical insights we have garnered so far. Briefly, the model relies on synaptic plasticity in the velocity-to-rotation ring connections and its mechanistic prerequisite (a rate code for the positional error instantiated in the rotation rings as shown in Fig. 4B) to achieve gain recalibration. We chose this mechanism, instead of other theoretical candidate mechanisms examined in the previous section, partly because we found it relatively easy to implement compared to others, but we conjecture that some of the other candidate mechanisms also have biologically feasible implementations. Thus, our model should be viewed as an example that demonstrates the effectiveness of our theoretical analysis rather than the only network model that can achieve gain recalibration. In addition to gain recalibration, the model can also reproduce two other important aspects of biological path integration: (1) flexible association of visual landmarks to different positions in the neural space and (2) correction of accumulated PI errors by visual landmarks. In the next subsections, we describe the model in detail and explain the mechanisms by which it achieves these aspects.

### 4.1 A connectivity pattern yielding a rate code for error

We begin by proposing a connectivity pattern that causes a neuron population to vary its firing rate monotonically with the difference in the bump locations of two other populations. As will be evident, this connectivity pattern plays a crucial role by providing the means to achieve a rate code for the positional error, with the error being the difference in the bump locations of the visual drive and the attractor activity.

The connectivity pattern can be described by considering three distinct neuron populations (X1-3 in Fig. 5A), each arranged on a circle like the populations in the ring attractor network. Suppose that populations X1 and X2 consist of excitatory and inhibitory neurons, respectively, and maintain their own activity bumps. The population X3 derives its activity based on the inputs from X1 and X2. The excitatory inputs from X1 to X3 are routed through topographic connections that wire together the neurons at the same angular location in the neural space. The inhibitory inputs from X2 to X3, however, are routed through CCW *offset* connections that wire together the neurons at different angular locations. To understand how this connectivity causes X3 to vary its firing rate as a monotonic function of the difference in the bump locations of X1 and X2, we can track the flow of neural activity as follows:

**Figure 5:**
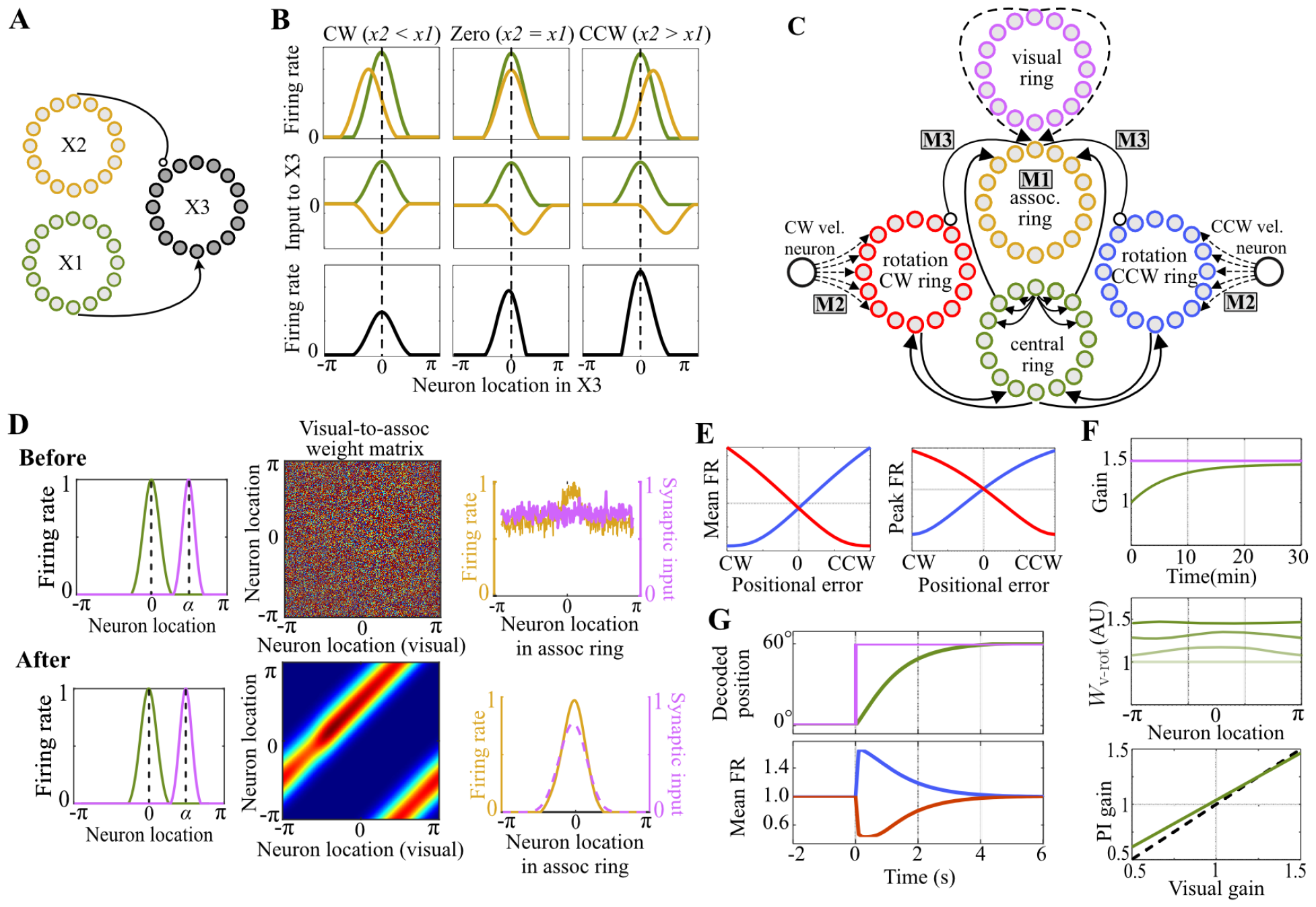
A modified ring attractor network model. (A) A proposed connectivity pattern that varies firing rate of population X3 as a monotonic function of the difference in the activity-bump locations of populations X1 and X2. Arrow and circle terminals denote excitatory and inhibitory connections, respectively. (B) Schematic diagrams depicting the computation within the proposed connectivity in panel (A). The first row shows the activity of the X2 population (yellow) relative to the activity of the X1 population (green) for different conditions: CW difference (left column), no difference (middle column), and CCW difference (right column). The second row shows the synaptic inputs to the X3 population from the excitatory X1 (green line) and inhibitory X2 (yellow line) populations. The fourth row shows the resulting firing rates of the X3 population. (C) Schematic representation of the model. Solid and dashed lines denote hardwired and plastic connections, respectively. The labels M1, M2, M3, correspond to the three modifications made to the classical ring attractor: M1 is the association ring, M2 refers to the plasticity of the velocity-to-rotation-ring connections, and M3 refers to the hardwired, offset association-to-rotation connections. (D) Numerical simulation demonstrating the association of the visual ring’s activity with the central ring’s activity through Hebbian plasticity. Top and bottom rows show the initial and final values of the simulated variables. The left column shows the firing rates of the central (green) and visual (pink) rings. The middle column visualizes the weight matrix describing the visual-to-association ring connections. The right column shows the firing rates of the association ring (yellow) and the synaptic inputs from the visual ring (pink). (E) Tuning curves depicting the relationships between the rotation rings’ mean and peak firing rates vs. the error (left and right graph, respectively). The color coding is the same as panel C. (F) Numerical simulations of the gain recalibration within the proposed model. The top shows the the recalibration of PI gain (green) toward the visual gain (pink) in a selected simulation. The middle shows the progression of the weights of the velocity-to-rotation ring connections (four samples normalized to initial condition of the weights with the opacity changes from the lightest (*t* = 0 min) and to the darkest green (*t* = 30 min) corresponding to chronological order of the samples.). The bottom shows the final values of the PI gain for various visual gains (green line) and the hypothetical perfect recalibration (dashed black line). (G) Numerical simulation demonstrating how visual landmarks correct positional error. The top panel shows the progression of the bump locations of the visual (pink) and central rings (green). Bottom panel shows the mean firing rate of CW (red) and CCW (blue) rotation rings over time.

Let *x*_1_ and *x*_2_ denote the location of X1 and X2’s activity bumps on the circular neural space, respectively. When *x*_2_ *− x*_1_ = 0, these activities are aligned, but the synaptic inputs to the population X3 from X1 and X2 are misaligned due to the CCW offset in X2-to-X3 connections (the second column of Fig. 5B). When *x*_2_ *−x*_1_ ≠ 0, however, X1 and X2’s activity bumps are misaligned with a CW (*x*_2_ *−x*_1_ *<* 0) or CCW (*x*_2_ *−x*_1_ *>* 0) difference in their locations. Consider first the CW-difference case (the first column of Fig. 5B). In this case, the bump location of X2 is shifted in the CW direction compared to that of X1. Because of the CCW offset in the X2-to-X3 connections, this misalignment between the activity bumps decreases at the level of synaptic inputs, bringing the inhibition from X1 closer to active neurons of X3, thereby decreasing the firing rate of X3 compared to the no-difference case. Consider next the CCW-difference case (the third column of Fig. 5C). In this case, the CCW offset in the X2-to-X3 connections redirects the inhibition from X2 further away from the active neurons of X3, thus increasing the firing rate of X3. Through this mechanism, the proposed connectivity pattern causes population X3 to vary its firing rate as a monotonically increasing function of the difference in the bump locations of X1 and X2. The direction of this monotonic relationship can be easily reversed if the offset in the X2-to-X3 connections is reversed from CCW to CW.

How can we make use of this connectivity pattern in our modified ring attractor network that will rely on plastic velocity-to-rotation ring connections for gain recalibration? The connectivity lends itself naturally to this recalibration mechanism, as it requires the rotation rings (X3) to vary their firing rates monotonically with the positional error, a quantity equal to the difference in the bump locations of the central ring (X1) and the visual drive (X2). Despite this suitability, however, employing the connectivity pattern in the ring attractor network requires an additional modification for a reason which will be clear in the next section.

### 4.2 Flexible Association of Landmarks to Positions through Hebbian Plasticity

In traditional ring attractor models, feedback from landmarks is incorporated into the network via direct synaptic connections from the visual ring [20, 46, 54]. In these connections, synaptic weights between coactive neuron pairs encoding the same position of the animal are potentiated through Hebbian plasticity, resulting in a flexible associative mapping between the visual ring and the attractor network [20, 55].

However, this approach, relying on direct plastic connections from the visual ring onto the attractor network, is incompatible with the connectivity pattern proposed in the previous section for the error-rate code. That is, the error code’s connectivity pattern requires the wiring of neurons representing different positions by an *offset* as opposed to Hebbian plasticity wiring together the neurons representing the same position. To resolve this incompatibility, our modified ring attractor network model makes a small change to the approach in traditional models by placing the plastic associative mapping problem outside the network, which makes it possible to include the error code’s connectivity pattern inside the network.

The modification is as follows: We first remove the visual-to-central ring connections (➄ in Fig. 1A) and introduce an intermediate ring of neurons, which we call an *association ring*, that associates the activity in the visual ring with that in the central ring by receiving inputs from both (see M1 in Fig. 5C). The afferent connections to this ring from the central ring are hardwired and weak, while those from the visual ring are plastic, hence capable of becoming strong. In a novel environment, where the plastic visual connections are initially untuned and random, the spatial selectivity of the visual ring’s activity is not conveyed in its synaptic inputs to the association ring. In contrast, inputs from the central ring to the association ring always preserve the spatial selectivity of the central ring’s activity because of the hard-wired, topographic connections. This combination of malleable inputs from the visual ring and weak but hardwired inputs from the central ring biases the visual-to-association ring connections near the central ring’s activity bump to be selectively potentiated through Hebbian plasticity (top row in Fig. 5D). Eventually, synaptic inputs from the visual ring become sufficiently strong and aligned with the inputs from the central ring, making the association ring’s activity strongly visually driven such that it implicitly represents a flexible associative mapping between the representations in the visual ring and the attractor network (bottom row Fig. 5D).

Thus, by serving as an intermediary, the association ring promises to act in our modified ring attractor network model as a proxy visual drive that can circumvent the previously noted incompatibility issues with the error-rate code’s connectivity pattern. In the next section, we describe how the association ring can be combined with this connectivity pattern to implement gain recalibration.

### 4.3 Gain Recalibration by Landmarks through Hebbian Plasticity

As noted earlier, our ring attractor network model relies on plasticity in the velocity-to-rotation ring connections for gain recalibration (M2 in Fig. 5C). However, this recalibration mechanism requires as its mechanistic prerequisite (described in Section 3.3) that CCW and CW rotation rings respectively increase and decrease their firing rates with the positional error in the central ring’s representation relative to the visual landmarks. To obtain these error-rate codes, we make use of the visual information in the association ring by connecting it to the CCW and CW rotation rings with CCW and CW *offset*, respectively (M3 in Fig. 5C). Combined with the topographic central-to-rotation ring connections, these *offset* connections from the association ring implement the connectivity pattern described in Section 4.1, thereby achieving the required error-rate codes (Fig. 5E). With these error-rate codes in the rotation rings and the plasticity in the velocity-to-rotation ring connection, our ring attractor model now includes all the necessary ingredients for gain recalibration.

To test if these ingredients are also sufficient to achieve gain recalibration, we first take an analytical approach based on Equation (15), which states a sufficient condition for gain recalibration. According to this condition, gain recalibration is guaranteed if the temporal change in the gain’s spatial average *k*_0_ has a positive slope with respect to the product of the animal’s velocity *v* and the attractor’s positional error 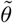. As discussed in Section 3.3.1, the gain’s spatial average *k*_0_ is positively correlated with the average strength of velocity-to-rotation ring connections. Therefore, the sufficient condition can be rephrased as follows: the gain recalibration is guaranteed if the temporal change in the average strength of velocity-to-rotation ring connections has a positive slope with respect to the product of *v* and 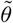. The presynaptic side of these plastic connections has velocity neurons encoding the animal’s velocity *v*, while the postsynaptic side has the rotation ring neurons encoding the positional error 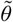. Since both of these rate codes have the same slopes on the CW and CCW parts of the network (negative on the CW part, positive on the CCW part), their correlated activity, as the driver of Hebbian plasticity, modifies the strength of velocity-to-rotation ring connections with a positive slope relative to the product of *v* and 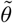, satisfying the sufficient condition in (15). However, as in Example 2 in Section 3.2.2, gain recalibration is expected to be imperfect due to imperfect encoding of the positional error 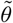 in the rotation rings whose firing rates are additionally modulated by the animal’s velocity *v*. As a result, our model is expected to recalibrate its average PI gain *k*_0_ to steady-state values that are close to, but not the same as, the visual gain, *k*^⋆^. During this recalibration, the model is expected to go through a transitory stage with a spatially non-homogenous PI gain *k*(*θ*) as Hebbian plasticity modifies the spatially distributed velocity-to-rotation ring connections non-uniformly because of non-uniform firing rate of rotation rings’ neurons across the neural space at any moment in time. However, as the animal runs in the environment, these non-uniform effects will be washed out, resulting in a spatially homogeneous PI gain *k*(*θ*). Mathematical details of these analytical findings are given in Appendix 6.5.

We next verify these analytical findings by performing numerical simulations of our modified ring attractor model for a simulated rat running on a circular track while visual landmarks were moved as per the visual gain *k*^⋆^. The model demonstrated imperfect yet stable gain recalibration for a range of *k*^⋆^ values (Fig. 5F).

### 4.4 Correction of Positional Errors by Landmarks through a Rate Code of Error

Next, we test if visual landmarks can correct PI errors in our model. The PI error manifests itself as a misalignment between the peak locations of the path-integration driven activity bump in the central ring and the strongly visually driven activity bump in the association ring. Traditional models correct for this misalignment by providing visual drive *directly* onto the central ring (like in Fig. 1A), toward which its activity bump is gravitated by means of attractor dynamics. In our model, however, we have removed such direct connections in Section 4.2 because of their incompatibility with the connectivity pattern needed for the error-rate codes. Instead, we have connected the association ring to the rotation rings with some offset. The question then arises: are these offset connections sufficient for PI error correction or do we need to reinstate direct connections onto the central ring from the visual drive?

As explained in the previous section, the offset connections causes the CCW and CW rotation rings to increase and decrease their firing rates monotonically with the positional error, respectively. This differential modulation of the rotation rings’ firing rates by the error is similar to their differential modulation by the velocity; when the animal is moving, the firing rate of one rotation ring increases and that of the the other decreases, which in turn shifts the activity bump along the central ring. Therefore, by employing rate codes of the positional error, our model effectively transforms the positional error into a virtual velocity signal, shifting the activity bump along the central ring in a manner decreasing this error—a manifestation of error correction provided by visual landmarks. We verified this error correction mechanism in numerical simulation of our model. Following a positional error introduced abruptly between the activity bumps of the ring attractor and the association ring, the differential changes in the rotation rings’ firing rates eliminated the error by re-aligning the central ring’s activity bump with that of the association ring (Fig. 5E).

## 5 Discussion

Fine-tuning of a neural integration computation is crucial to maintain accurate representations of continuous variables since the relationship between the sensing of the relative change in a continuous variable and its actual value can fluctuate on both developmental (e.g., changes in body size that can affect location coding) and behavioral timescales (e.g., swimming versus walking in the case of location coding) and even due to dynamic biological processes, such as circadian rhythm, that can alter synaptic transmission and intrinsic electrical properties of neurons. Building upon previous behavioral work on perceptual plasticity of human locomotion [56], physiological evidence for such fine-tuning was first observed in hippocampal place cells [35], where persistent conflict between self-motion and external visual cues re-calibrated the integrator gain. In the present paper, we give the first theoretical examination of this phenomenon in continuous bump attractor networks (CBANs), a prevailing model for representations of continuous variables.

Our examination unveiled the algorithmic and mechanistic requirements for gain recalibration in a ring attractor network, a representative CBAN model used for circular continuous variables. In CBAN models, when the integration gain is inaccurate, an internal representation of a continuous variable slightly drifts relative to its actual value, resulting in encoding errors. Absolute ‘ground-truth’ information, such as feedback from visual landmarks for location coding, correct these errors through internal dynamics of the network, without the need for an explicit rate-based representation of the error. In contrast to this automatic error correction through network dynamics, we found that fine-tuning the integration gain based on errors requires an *explicit error signal*, i.e., firing rate of some neurons to encode the error in the CBAN’s representation of a continuous variable relative to its actual value. Building upon this insight, we also proposed a ring attractor network model that shows how a CBAN can recalibrate its integration gain through biologically known plasticity mechanisms. Although the ring attractor is specialized for integration of 1D circular continuous variables (e.g., an animal’s location on a circular track), our findings can be readily extended to higher dimensions and other types of continuous variables. Overall, our findings suggest that a rate code for the error in the internal representation of a continuous variable is a core component of the bump attractor-type neural integrators, and that such a rate code plays an essential role in their gain recalibration.

### 5.1 The Bump Attractor Network as an Adaptive Kalman Filter

To identify algorithmic requirements for recalibration of the integration gain, we simplified dynamics of the ring attractor through a dimensionality reduction protocol described in [41]. Similar approaches have been successfully applied in recent years to explore the neural dynamics capturing how high-dimensional neural data evolves within low-dimensional topological structures [6, 14, 57]. In our specific case, the dimensionality reduction led to a simplified 1D model of the ring attractor, capturing the dynamics of its representation as a function of external inputs that provide differential (e.g., animal’s velocity) and absolute (e.g., positional feedback from visual landmarks) information [41, 42, 58].

Previous research showed that, when two external cues are presented as inputs, the ring attractor network fuses them optimally in the Bayesian sense [20, 59–61]. Furthermore, if one of the cues provides only differential information like the animal’s velocity that is integrated over time to compute the overall change in the continuous variable, the ring attractor network performs the Bayesian fusion recursively for each step of the integration [41]. This recursive computation is known as Kalman filtering and has been proposed as a model of cue integration in the entorhinal cortex of the mammalian brain [21]. Consistent with this prior work, we found that the ring attractor network operates as a Kalman filter updating the representation by a combination of integrated relative information (the internal model component of the Kalman filter) and the instantaneous feedback from absolute information (the measurement model component).

In engineered systems, the accuracy of a Kalman filter relies on precise knowledge of its internal model parameters; to address this issue, adaptive Kalman filters that fine-tune their own parameters have been proposed [62, 63]. Like adaptive Kalman filters, we showed that a ring attractor network can fine-tune itself through gain recalibration. We also elucidated the algorithmic requirement for this recalibration, showing that the integration gain must change in the same direction as the product of the animal’s velocity and the error in the attractor’s representation relative to the absolute ‘ground-truth’ information. Interestingly, this requirement resembles characteristics of a classical algorithm known as the MIT rule in adaptive control systems and Kalman filtering [64]. Thus, a ring attractor with gain recalibration effectively operates as an adaptive Kalman filter, updating its representation accurately through a finely tuned integration gain.

### 5.2 Necessity of a Rate-Based Explicit Error Signal in the Bump Attractor Networks

Satisfying the algorithmic requirement for gain recalibration is subject to certain mechanistic constraints, which we discovered analyzing the network dynamics. In essence, gain recalibration requires that some neurons vary their firing rates monotonically with the instantaneous value or the time-integral of the error in the representation of the continuous variable relative to its true value. Without such error-rate codes, the network does not have a teaching signal that can guide tuning of its gain for recalibration. This shows that, for CBAN networks, learning from representational errors to recalibrate the integration gain is a very different neural process than correcting the errors. In the case of error correction, input signals from absolute ‘ground-truth’ information sources, such as visual landmarks for a CBAN encoding location, are sufficient to trigger an error correction response from network dynamics. In contrast, recalibrating the integration gain based on representational error additionally requires a neural signal that explicitly encodes this error via a rate code.

This hypothesized error signal resembles reward and sensory prediction error signals within the mammalian brain. Dopamine neurons in the midbrain of mammals encode error in the internal predictions of reward via monotonic changes in their firing rates [65,66]; they exhibit elevated activity with more reward than predicted, remain at baseline activity for fully predicted rewards, and exhibit depressed activity with less reward than predicted. Climbing fiber inputs to Purkinje cells of the mammalian cerebellum encode errors in the predicted sensory consequences of motor commands relative to the actual sensory feedback via changes in the rate and duration of complex spikes [67, 68]. Both of these prediction error codes are thought to act as a teaching signal that fine-tunes the internal models, mapping the stimulus to reward prediction in the dopamine system and the motor commands to sensory prediction in the cerebellum, through plasticity, just like how error coding can act as a teaching signal that recalibrates the integration gain of a CBAN.

To instantiate this idea computationally, we presented a modified ring attractor model that can recalibrate its integrator gain based on a rate code of the instantaneous error via Hebbian plasticity. Relevant to this model, a previous study hypothesized a currently unknown plasticity rule as a mechanism for gain recalibration within the ring attractor network [32]; the hypothesized plasticity rule modifies synaptic weights of each neuron according to an implicit positional error signal, computed locally within each neuron through comparison of the synaptic inputs at its basal and apical dendrites receiving, respectively, the absolute ‘ground-truth’ information and the network’s current representation information. While it is unclear whether such a plasticity rule exists in the brain, our model demonstrates a biologically plausible alternative: Hebbian plasticity, combined with an explicit rate-based representation of the error, is sufficient to achieve gain recalibration. It remains as a future work to experimentally test if such an error signal exists in the brain circuits that are thought to employ CBANs for encoding continuous variables.

### 5.3 Implications of a Distributed, Inhomogenous Integration Gain

Prior CBAN models implicitly assumed the integration gain to be a single, global parameter of the network, independent of the value of the encoded continuous variable [11, 31]. Although the idea of different, hard-wired integration gains has previously been suggested in the context of location coding to explain the changes in the spatial scale of place coding along the dorsal-ventral axis of the hippocampus [27], it is assumed that the integration gains are constant at all locations within an environment. In contrast, we showed that the integration gain within a single CBAN, specifically the ring-attractor network, is a distributed parameter instantiated in the network’s array of synaptic weights, implying that the network can adopt unique integration gains for different values of the encoded continuous variable. This would be, for example, a CBAN changing its integration gain depending on the location of the animal in the context of spatial navigation or depending on the amount of accumulated evidence in the context of decision-making. In the latter case, the CBAN may look like it unevenly weights early or late evidence, which is a well-established phenomenon known as primacy and recency effects in the decision-making literature [69–71]. Compared to a network with a single, global integration gain, a network with a distributed, possibly inhomogeneous, gain can adjust its representation metric for the continuous variable locally, hence providing greater flexibility in representing different values of the continuous variable with uneven resolutions, depending on, for instance, their behavioral significance [72].

How might gain inhomogenoity arise? Theoretically, it can be a product of the recalibration process if the teaching signal (e.g., feedback from absolute ‘ground-truth’ information sources) is available unevenly across the values of the continuous variable. In the context of location coding, for example, such differences may occur between when the animal is nearby the boundaries of the environment, which is a relatively rich area in terms of external ground-truth information, versus when it is near the center of the arena. We speculate that such spatially distributed recalibration of the integration gain may offer a mechanistic explanation to some of the experimental findings about local distortions and deformations in the activity patterns of entorhinal grid cells and hippocampal place cells, encoding the animal’s location, during environmental manipulations [73–80]. According to this speculation, grid patterns might get distorted nearby environmental boundaries through changes in the local integration gain of the network; through the same mechanism, place cells might represent locations nearby landmarks and boundaries with a greater spatial resolution (also known as overrepresentation). Overall, inhomogenous integration gain of CBANs offer a potential explanation to an array of seemingly complex responses in spatial navigation as well as other brain functions.

## 6 Appendix

### 6.1 Model Setup: Ring attractor network

Dynamics of the ring attractor network can be formulated by modeling the firing rate of each neuron in response to its synaptic inputs.

#### 6.1.1 Dynamics of the central ring

We begin with the central ring. Parameterizing neurons based on their locations *ψ ∈ S*^1^ in the central ring, we can model their dynamics via

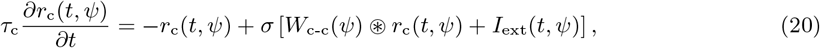

where *r*_c_(*t, ψ*) denotes the firing rate of the central ring neuron *ψ* at time *t, τ*_c_ denotes the synaptic time constant, ⊛ denotes the circular convolution operation, *σ* is an activation function (chosen in the present work as a rectified linear unit, RELU), and *W*_c-c_ : *S*^1^ *→* ℝ is a rotationally invariant synaptic weight function (chosen in the present work as a symmetric local excitation global inhibition, LEGI), and *I*_ext_(*t, ψ*) denotes the external synaptic input to the central ring neuron *ψ* at time *t* from CW, CCW rotation rings and the visual ring.

We can write this combined external input to the central ring as

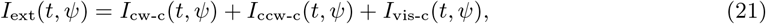

where *I*_cw-c_, *I*_ccw-c_, and *I*_vis-c_ denote the inputs to the central ring from CW, CCW, and visual rings.^1^ Using synaptic weights and firing rates, we can compute each of these inputs according to the equation

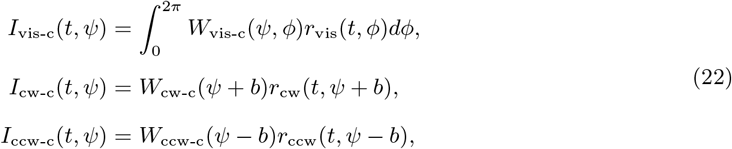

where *W*_vis-c_ : *S*^2^ *→* ℝ denotes the synaptic weight function describing the all-to-all connections between the visual ring neurons and the central ring neurons, *r*_vis_ denotes the firing rate of visual ring neurons, *W*_cw-c_, *W*_ccw-c_ : *S*^1^ *→* ℝ _*≥*0_ denotes the synaptic weight functions describing the one-to-one and differentially offset connections between the CW,CCW rotation ring neurons and the central ring neurons, and *b ∈* ℝ denotes the amount of this connection offset. Collectively, Equations (20)–(22) constitute the central ring model considered in this paper.

#### 6.1.2 Dynamics of the rotation rings

We next model firing rates of the rotation rings’ neurons. Similar to previous work [81], we assume that synaptic time constants of the rotation rings are much faster than that of the central ring. This allows us to model the firing rates of rotation rings’ neurons as algebraic quantities whose values are determined instantaneously by the synaptic inputs they receive, namely,

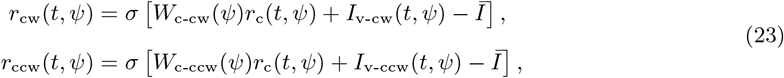

where *W*_c-cw_, *W*_c-ccw_ : *S*^1^ *→* ℝ _*≥*0_ denote the synaptic weights describing the one-to-one connections between the central ring and the each rotation ring, *Ī* denotes global inhibition, and *I*_v-cw_, *I*_v-ccw_ denote the synaptic inputs from the CW, CCW velocity neurons to the respective rotation rings, namely

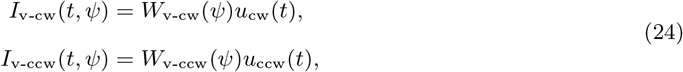

where *W*_v-cw_, *W*_v-ccw_ : *S*^1^ *→* ℝ _*≥*0_ denote the synaptic weight functions describing the one-to-all connections between the velocity neurons and the respective rotation rings, and *u*_cw_, *u*_ccw_ denotes the firing rates of the velocity neurons. Following previous work [31], we assume that these firing rates simply follow a linear relationship with the animal’s velocity *v* as follows:

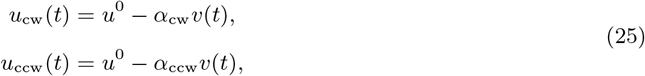

where *u*^0^ *≥* 0 denote the intercept of velocity neurons’ tuning curves, and *α*_cw_ *>* 0, *α*_ccw_ *>* 0 denote the slopes of the velocity neurons’ tuning curves. Substituting this model of velocity neurons into Equation (24), we obtain a more explicit expression for the synaptic inputs from the velocity neurons to the rotation rings as follows:

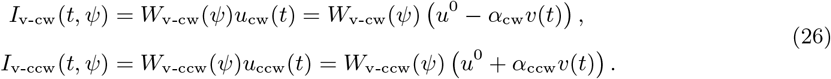

Collectively, Equations (23), and (26) constitute the model of the rotation rings.

### 6.2 Model reduction of network dynamics

#### 6.2.1 An ansatz solution to the central ring

It is well known that when the recurrent weight function *W*_c-c_ has the classical pattern of local excitation and global inhibition (LEGI), a persistent activity bump emerges in the central ring [1,38–40]. As done in the previous work [41, 42], this bump of activity can be identified with a one-dimensional ansatz solution to Equation (20):

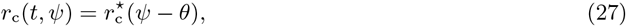

where *θ* denotes the location of the activity bump according to the ansatz function 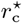 whose properties are stated in the Assumption below.

##### Assumption 1.

*The function* 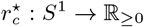 *has even symmetry and compact support around its single peak ψ* = *θ, namely*,

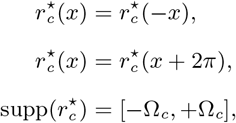

*where* Ω_*c*_ *denotes the limits of the support, corresponding to the half-width of the activity bump. Furthermore, the function* 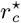 *is twice differentiable, and there exists a non-empty open neighborhood N of the origin where the second derivative of* 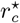 *is negative, namely*,

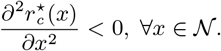

Δ

The LEGI pattern of recurrent weight function *W*_c-c_ of the central ring maintains the shape of the activity bump 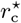, but does not control the location *θ* of the bump over the neural space. Rather, the external inputs *I*_ext_ to the central ring control *θ*. Therefore, for a given smoothly varying, external input *I*_ext_*t, ψ*, the ansatz solution 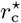 to Equation (20) satisfies

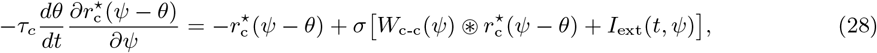

since one can show (using the chain rule) that 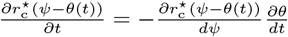.

#### 6.2.2 An ansatz solution to the rotation rings

Substituting the ansatz solution 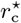 for the central ring dynamics into Equations (23) and (26), we can also solve for the firing rates of the rotation rings as follows:

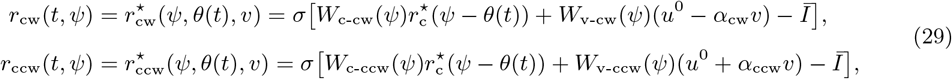

where we introduced the functions 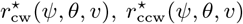 as a shorthand notation for the ansatz solutions of the activity of the rotation rings. Since firing rates of rotation rings’ neurons depend on the ansatz solution 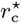 for the central ring, the ansatz solutions 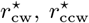 for the rotation rings can have a compact support interval like 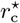, as formally stated below:

##### Assumption 2.

*The global inhibition Ī to the CW and CCW rotation rings is sufficiently high such that*

1. *for a given velocity v and the bump location θ, the functions* 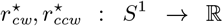 *have compact support intervals over ψ:*

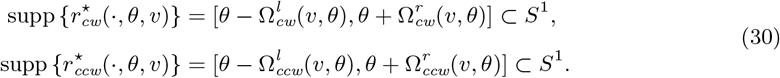

*where the half widths* 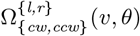 *potentially depend on velocity v and bump location θ. We also assume the total widths are not zero and do not cover the entire extent of S*^1^, *namely*,

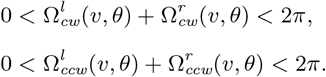
2. *2 these compact support intervals are contained inside 𝒩, the open set where the second derivative of the ansatz function* 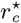 *is negative as described in Assumption 1. △*

Notice that we assume the support intervals of the rotation rings’ persistent activity bumps 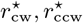 during the animal’s immobility are defined relative to the peak location *θ* of the central ring’s activity bump 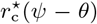. However, unlike the central ring’s activity bump, the widths of the rotation rings’ activity bumps are not necessarily fixed. That is, the left and right half-widths of the activity bumps may vary with *v* and *θ*, depending on the animal’s velocity and the spatial variance of the synaptic weight functions *W*_c-cw_, *W*_c-ccw_, *W*_v-cw_, and *W*_v-ccw_.

#### 6.2.3 Dynamics of the position representation *θ* during animal’s immobility in the absence of landmarks

During the animal’s immobility (i.e., *v* = 0), Equation (29), the solution to rotation ring dynamics, reduces to

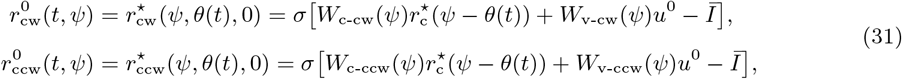

where the superscript 0 denotes that a quantity is evaluated when the animal is stationary (i.e., *v* = 0) without visual landmarks. Furthermore, when there are no landmarks, the firing rate of visual ring neurons becomes 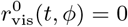 for all *t, ϕ*. Substituting these solutions into Equation (22), we obtain

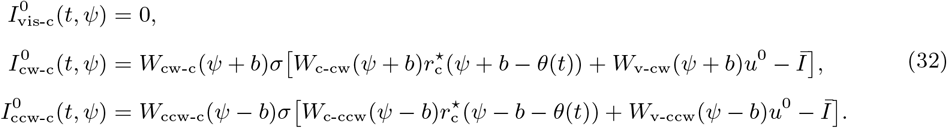

Combining these synaptic inputs according to Equation (21), we obtain the total external input to the central ring during the animal’s immobility in the absence of landmarks as

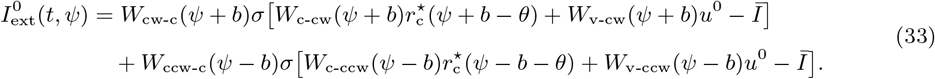

The external synaptic input *I*_ext_ drives the temporal changes in *θ* according to the differential equation in (28). In a stable implementation of the path integration (PI), 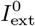 must be such that the equilibrium conditions of that differential equation is satisfied as shown below:

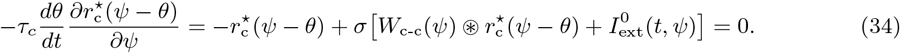

This implies that *θ* remains constant during immobility of the animal, namely,

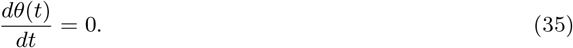

In the remark below, we describe a necessary property of 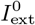to satisfy these equilibrium conditions.

##### Remark 1.

*In steady-state, Equation (34) requires the terms on its right-hand side to satisfy*

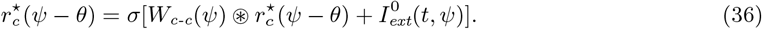

*This requirement can be translated into different requirements for the cases that ψ is inside or outside the support interval of* 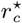 *as follows:*

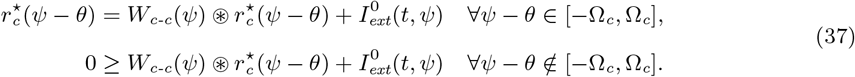

*Since* 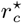 *and W*_*c-c*_ *are even symmetric functions about the origin, the first line in Equation* (37) *implies that the total external synaptic input* 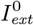 *must also be even symmetric inside the interval* [*−*Ω_*c*_, Ω_*c*_] *independent of the choice of b, namely*,

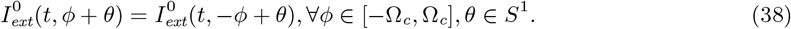

Δ

Note that in the present work, we constrain the model parameters such that this symmetry property of 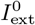, is satisfied to ensure stable PI.

#### 6.2.4 Dynamics of the position representation *θ* without visual landmarks but with movement, i.e. pure path integration

In the present section, we follow the dimensionality reduction protocol described in [41] to derive the one-dimensional differential equation model of the position representation *θ* as a function of the animal’s velocity *v*. Briefly, this protocol first computes the change in the external synaptic input of the central ring with the movement compared to the baseline case of animal’s immobility, and then map this input change to a change in the position representation *θ*.

##### Computing the change in the synaptic inputs to the central ring with the movement

When the animal is immobile, CW and CCW velocity neurons fire at their baseline rates *u*_cw_ = *u*^0^ and *u*_ccw_ = *u*^0^. During movement of the animal, firing rates of these velocity neurons change in different directions:

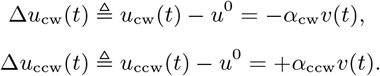

These directionally different changes in the firing rates of the velocity neurons carry over into their synaptic drive to the rotation rings via

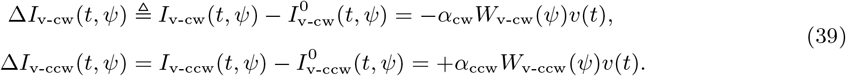

Because the firing rates of the rotation ring neurons are determined instantaneously by their synaptic inputs, these input changes during the movement lead to instantaneous perturbation of their activity bumps during the immobility. To find an approximate analytical formulation of these perturbations, we follow the previous work [41] and linearize the algebraic model of rotation ring firing rates, given in Equation (23), about the immobility condition *v* = 0:

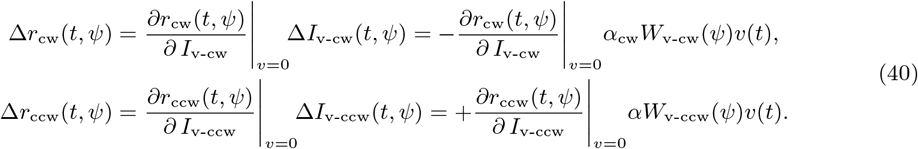

Explicitly differentiating the algebraic model (23), we obtain

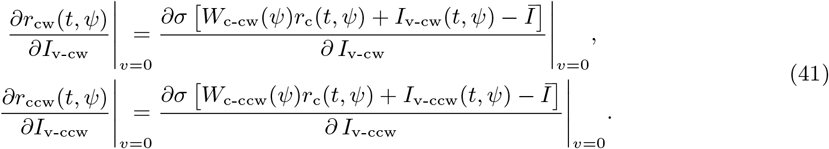

To find closed-form, analytical expressions of these terms, we follow the previous work [41] that approximates the derivative of the linear threshold-type activation function *σ* as follows:

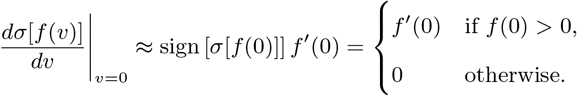

Using this approximation and Equation (23), we obtain

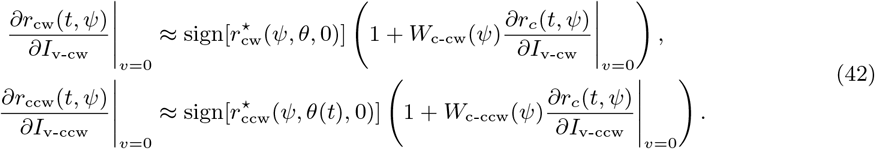

Recall that firing rate of the central ring neurons change according the differential equation in (20), implying that they are independent of the received synaptic inputs at a given time, namely,

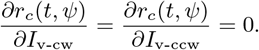

Substituting this into Equation (42) leads to

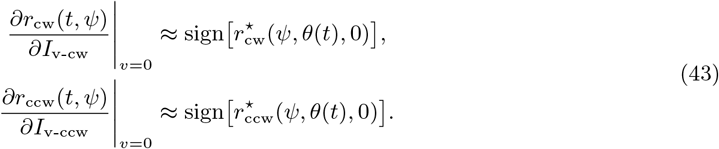

Substituting these expressions into Equation (40), we finally obtain closed-form, analytical expressions of the firing rate perturbations for rotation rings as

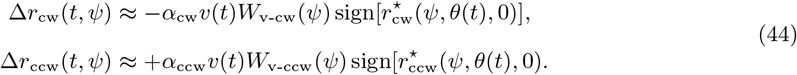

To map these firing rate perturbations into the disturbances in the synaptic inputs to the central ring, we return to Equation (22). First, we substitute 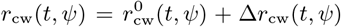 into the first line of Equation (22) and 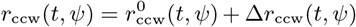 into the second line. After subtracting the results of these substitutions from the last two lines in Equation (32), we obtain

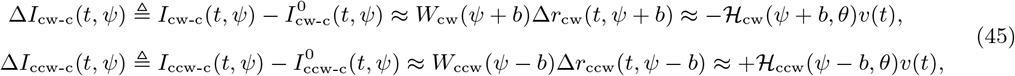

where

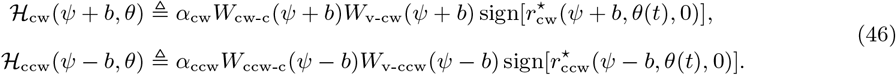

We can compute the total disturbance by simply adding up these individual disturbances:

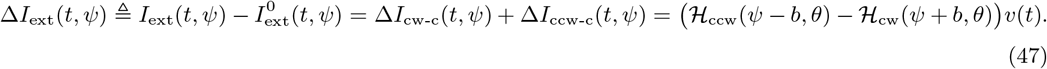

##### Relation between the synaptic input change and the change in *θ*

We next derive how this synaptic input disturbance modifies the location *θ* of the activity bump, described by the ansatz function 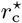. For this derivation, we substitute the above equation for the synaptic input disturbance into Equation (28), namely the dynamics of *θ*. This substitution yields

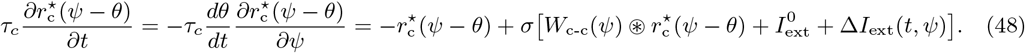

Recall that, when Δ*I*_ext_(*t, ψ*) = 0 and 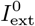 satisfies the symmetry requirement in Equation (38), the above differential equation reduces to the equilibrium condition in Equation (35), namely,

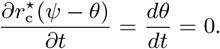

Thus, the addition of dynamic synaptic input Δ*I*_ext_(*t, ψ*) into Equation (48) perturbs the dynamics away from these equilibrium conditions.

As previously assumed, the attractor dynamics allow only rotation of the bump (i.e., change in *θ*) in response to external inputs. To find the resulting rotation in response to the disturbance Δ*I*_ext_, we first project^2^ Δ*I*_ext_ onto the tangent subspace of the ansatz function 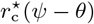, described by the gradient of 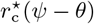 with respect to *θ*, as follows:

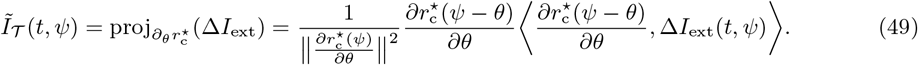

We then replace the term Δ*I*_ext_ in the central ring dynamics in Equation (48) with its tangential component *Ĩ*_*𝒯*_, which leads to

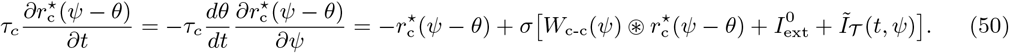

Since the right hand side of this equation is zero when *Ĩ* _*𝒯*_ = 0 (see Equation (34)), we can reduce the right-hand side to

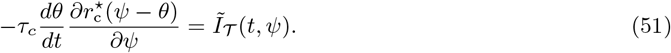

See the remarks and the proposition at the end of this section for a formal proof of this reduction. Using Equation (49), we can rewrite this differential equation more explicitly as

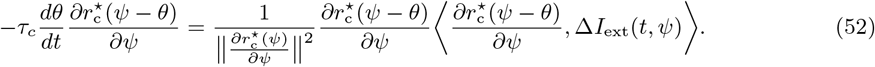

Eliminating the common terms in both sides simplifies this differential equation to

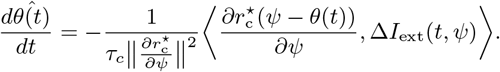

We then write the inner product more explicitly by using its integral-based definition, which yields

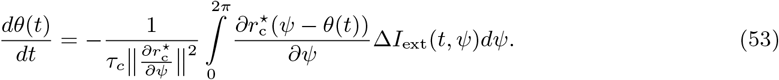

Since Δ*I*_ext_ depends linearly on the animal’s velocity *v* (see Equation (47)), this equation implies

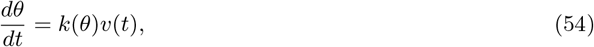

where *k* : *S*^1^ *→* ℝ_>0_ denotes the PI gain and takes the form

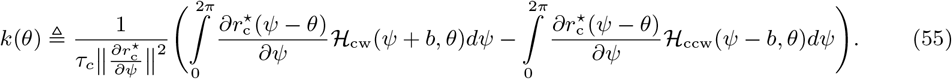

##### Recasting the analytical expression of the PI gain into a simpler form

Next, we manipulate Equation (55) to garner insight into the analytical expression of the PI gain and to simplify it. To this end, we begin by adding and subtracting the term (*ℋ*_ccw_(*ψ* + *b, θ*) *− ℋ*_cw_(*ψ* + *b, θ*)) */*2 to and from the first integrand and the term (*ℋ*_cw_(*ψ − b, θ*) *− ℋ*_ccw_(*ψ − b, θ)*) */*2 to and from the second integrand, which yields

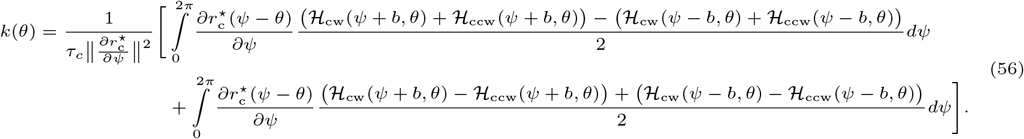

Equation (56) reveals that two integral terms contribute to the PI gain:

- The first integral quantifies the contribution of the *differential* component of the positively and negatively offset (*b*) inputs to the central ring from the rotation rings.
- The second integral quantifies the contribution of the residual, *common-mode* component of the positively and negatively offset (*b*) inputs to the central ring from the rotation rings.

As shown by the motivating examples in Fig. 6, the *differential* component is an odd function while the *common-mode* component is an even function. The integrals in Equation (56) compute the inner product of these components with an odd function, namely, the gradient of the ansatz 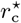. Thus, the first integral, including the *differential* component, yields a nonzero value but the second integral, including the *common-mode* component, yields zero. Based on these motivating examples, we assume that the second integral in Equation (56) has none or negligible effects on the gain, compared to the first integral.

**Figure 6:**
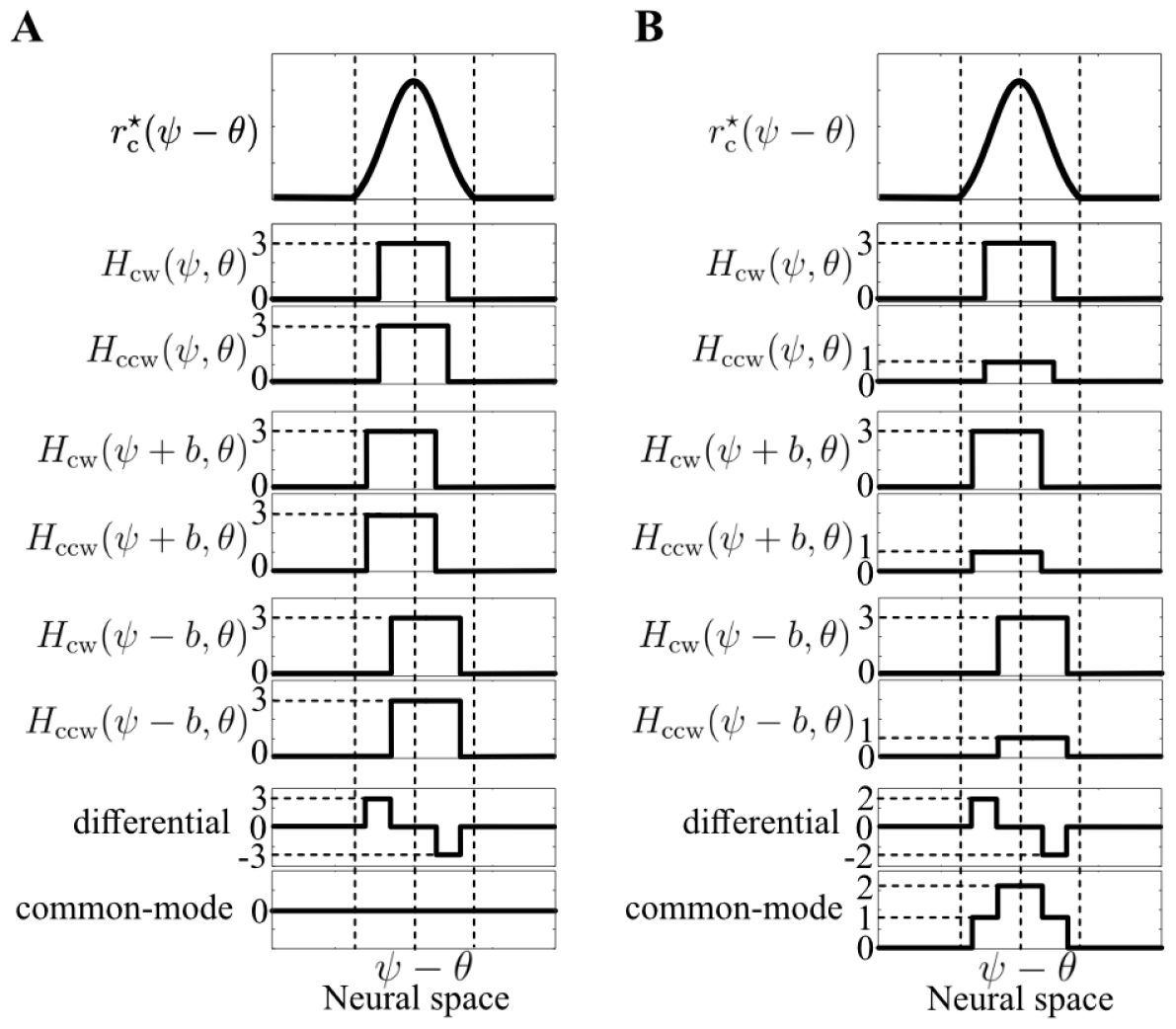
Examples showing that *common-mode* component results in an even function. (A) If the terms *H*_cw_,*H*_ccw_ are symmetric, the *common-mode* becomes zero, making the result of the second integral in Equation (56) also zero. (B) If the terms *H*_cw_,*H*_ccw_ are not symmetric, the *common-mode* results in a nonzero even function. Since the second-integral in Equation (56) is the inner product of the *common-mode* with an odd function, namely, the gradient of the ansatz 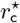, the result of the second integral again becomes zero.

###### Assumption 3.

*We assume that parameters of the ring attractor network satisfy*

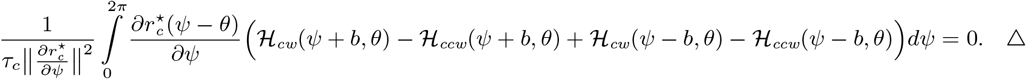

Under this assumption, Equation (56) reduces to

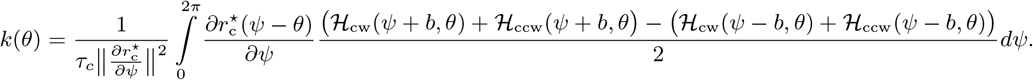

To further simplify this expression, we partition the integral into a difference of two integrals :

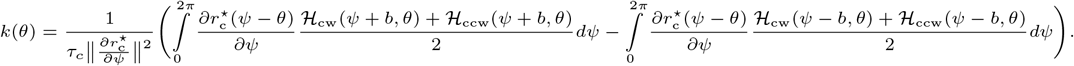

In this new form, we then perform a change of the integration variables to *z* ≜ *ψ − θ* + *b* for the first integral and to *z* ≜ *ψ − θ − b* for the second integral, which yields

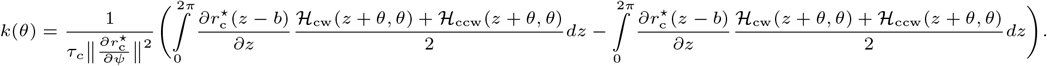

Note that when performing this change of variables, the integration bounds do not need to be updated since the integrands are periodic, and the bounds cover their entire period. These two integrals can thus be combined into a single integral as follows:

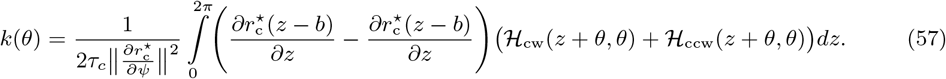

Assuming a sufficiently small offset *b*, we can employ the first-order Taylor series approximation to the difference term

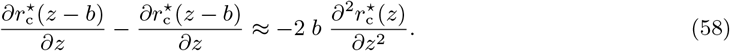

This approximation simplifies Equation (57), yielding a concise analytical expression of the PI gain

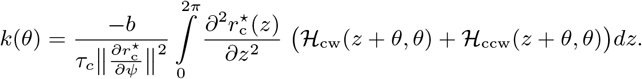

For notational consistency with the rest of the present paper, we make a change of variables once more and substitute *ψ* ≜ *z* + *θ* into this equation, which yields

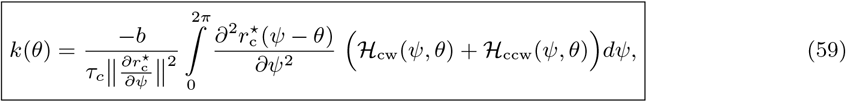

where the terms *H*_cw_,*H*_ccw_ are given as Equation (46). Throughout the paper, we mainly use this equation as an analytical expression for the PI gain. In the Remark below, we also would like to present an alternative expression that will be useful later when deriving mechanistic requirements for the gain recalibration.

###### Remark 2.

*To derive the alternative expression, we examine the support interval of the terms constituting the integral in Equation (59) and re-define the boundaries of the integral accordingly*.

*We begin this examination with the terms H*_*cw*_, *H*_*ccw*_. *By definition (see Equation (46)), these terms have the same compact support interval as the ansatz functions* 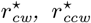, *describing the solution to activity of CW, CCW rotation ring during the animal’s immobility (v* = 0*), namely*,

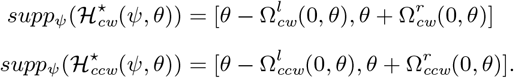

*Since union of compact sets is compact, support of the summation ℋ*_*cw*_ + *ℋ*_*ccw*_ *is also compact, i*.*e*.,

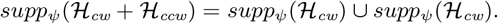

*Assuming that the union of the support intervals of ℋ*_*cw*_ *and ℋ*_*ccw*_ *is a connected set, we can express the support interval of the summation ℋ*_*cw*_ + *ℋ*_*ccw*_ *with a single interval as shown below:*

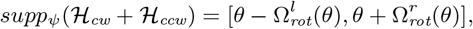

*where*

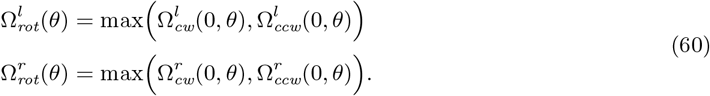

*We next examine the second derivative of the ansatz function* 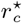. *Recall that, if a function has a compact support interval, then its derivatives enjoy the same interval. This implies*

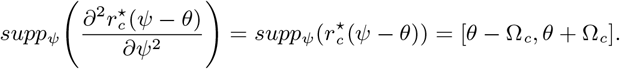

*It follows from the second part of Assumption 2 that the support of the summation term ℋ*_*cw*_ + *ℋ*_*ccw*_ *lies inside the support of the derivative term* 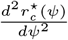 . *Therefore, the integrand term in Equation (59) attains the same compact support interval as the summation ℋ*_*cw*_ + *ℋ*_*ccw*_, *namely*,

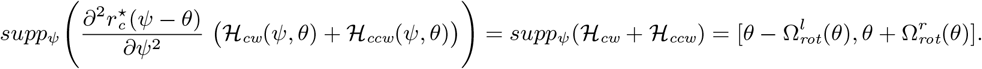

*This lets us reduce the bounds of the integration in Equation (59) to*

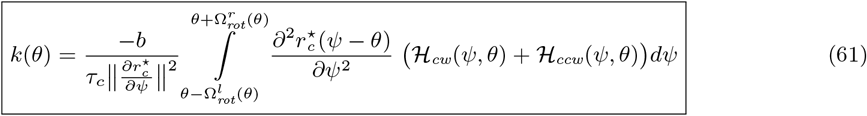

*Note that Equations (59) and (61) are analytically equivalent and used interchangeably throughout the paper as deemed appropriate. △*

Finally, we describe in the Remark below that similar steps can be followed to rewrite the equality in Assumption 3 in a more compact form.

###### Remark 3.

*By following the same steps as in between Equations (56)-(58), we obtain an alternate form for the equality in Assumption 3 as*

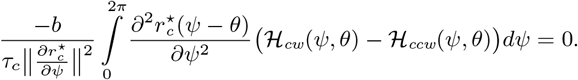

*Furthermore, by following the same steps as in Remark 2, we can choose the integration bounds more precisely as shown below:*

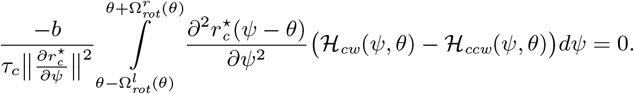

Δ

##### Steps to reduce right-hand side of Equation (50) to right-hand side of Equation (51)

###### Remark 4.

*As we mentioned previously, the ansatz function* 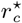 *and its derivative have the same compact support interval, namely*,

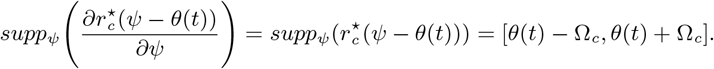

*Recall that, by definition in Equation (49), the tangential component Ĩ*_*𝒯*_ *of the disturbance* Δ*I*_*ext*_ *lies in the same direction as the derivative* 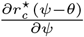. *This implies that*

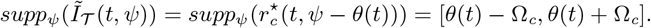

Δ

###### Remark 5.

*Let f* : *S*^1^ *→* [0, *∞*) *be a nonnegative function. Composition of the RELU activation function σ with f is equal to f, namely*,

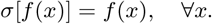

Δ

###### Assumption 4.

*Outside the support interval, the tangential component function Ĩ_𝒯_ and the ansatz function* 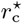 *are equal to zero, namely*,

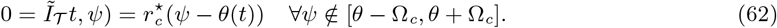

*Inside the support interval, they are not zero and assumed to satisfy*

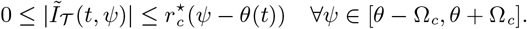

*Together, these two conditions imply*

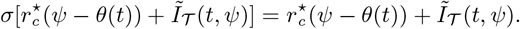

Δ

###### Proposition 1.

*The right-hand side of Equation (50) is equivalent to the right-hand side of Equation (51), namely*,

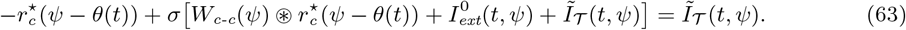

*Proof*. We prove the statement in the proposition by showing that Equation (63) is true in a case by case manner for *ψ* outside and inside the support interval of the ansatz function 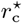.

1. Consider first that *ψ* is outside the support interval of the ansatz function 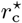. Since *Ĩ*_*𝒯*_ (*t, ψ*) is zero in this case as per Equation (62), Equation (63) reduces to

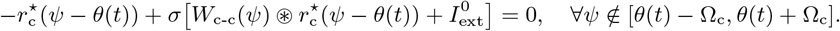

By Equation (34), this is true.
2. Consider now the remaining case that *ψ* is inside the support interval of the ansatz function 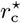. In this case, *Ĩ* (*t, ψ*) is not zero. Furthermore, as noted in Equation (37) the ansatz function satisfies

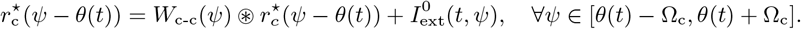

Using this equation, we can simplify Equation (63) to

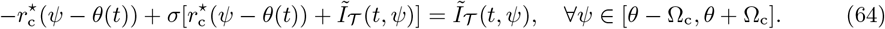

By Assumption 4, the activation function term on the left-hand side of this equation satisfies

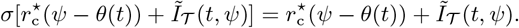

Based on this equation, we can verify that Equation (64) is true. □

#### 6.2.5 Dynamics of the position representation *θ* with visual landmarks

We continue following the dimensionality reduction protocol of [41] to extend Equation (54) of *θ* dynamics to the case where there is positional feedback from visual landmarks. To this end, we first extend the synaptic input disturbance Δ*I*_ext_ in Equation (47) by adding the visual component *I*_vis_, namely,

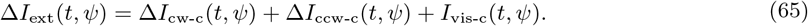

We then plug this extended disturbance into the differential equation in (53), which yields

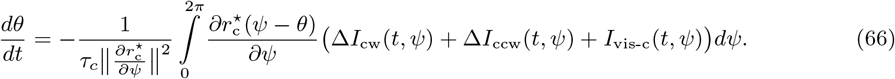

In the previous section, we analyzed this equation for *I*_vis-c_(*t, ψ*) = 0 and found

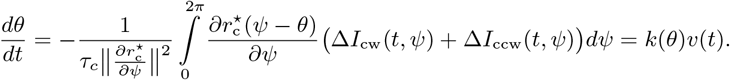

This lets us simplify Equation (66) to

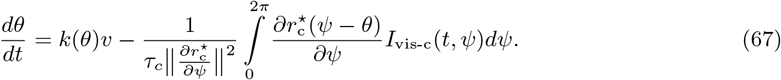

Next, we use Equation (7), expressing the assumed form of visual input Δ*I*_vis-c_, and obtain

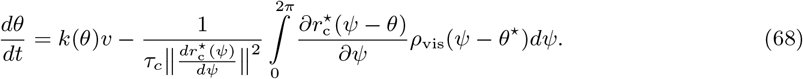

At the end of this section, we present three remarks. The first remark shows that the integral in Equation (68) can be reduced to a function, say *β* : *S*^1^ *→* ℝ, as follows:

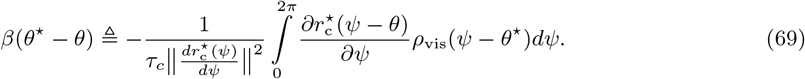

Substituting this function into Equation (68) yields a simple form for the differential equation of *θ* dynamics as follows:

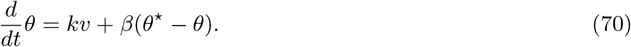

The second corollary shows that *β* is an odd function (i.e., *β*(*x*) = *−β*(*−x*)) because of even symmetry of the ansatz functions 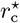 and *ρ*_vis_. Finally, note that, although we do not prove anything about the derivative of the function *β*, stable landmark control requires *β*^*′*^(0) *>* 0.

##### Remark 6.

*Let f and g be bounded continuous functions from S*^1^ *to* ℝ. *The dot product of f* (*x − y*_0_) *and g*(*x − z*_0_) *over x can be identified with a another function h* : *S*^1^ *→* ℝ *that depends on the difference y*_0_ *− z*_0_ *∈ S*^1^ *as shown below:*

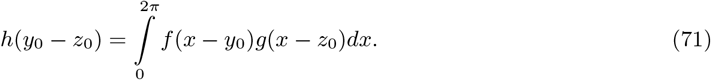

*To see this, let us expand f and g into their Fourier series representation*

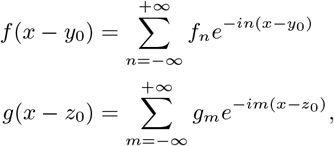

*where f*_n_ *and g*_m_ *denote Fourier series coefficient of the functions f and g, respectively. Using these expansions, we can compute the integral as follows:*

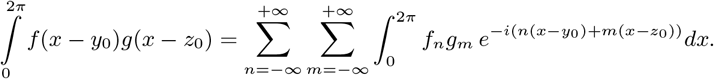

*When m ≠ − n, the integral vanishes. Thus, we can reduce the double summation to a single summation with m* = *−n and simplify the integral as follows:*

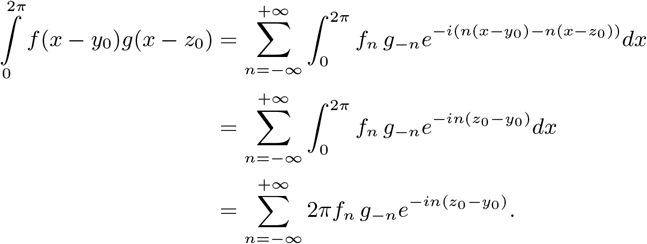

*The resulting summation can be identified as the Fourier series expansion of another function, say h* : 𝕊^1^ *→* ℝ, *such that*

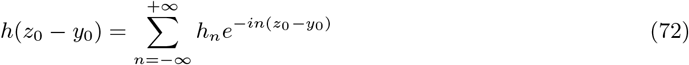

*where h*_n_ := 2*πf*_n_*g*_*−*n_ *denotes the Fourier series coefficients of the function h*. Δ

##### Remark 7.

*If bounded continuous functions f* : *S*^1^ *→* ℝ *and g* : *S*^1^ *→* ℝ *have even and odd symmetry about the origin, then their dot product, as defined in Equation (71) of the Remark 6, is an odd function, namely*,

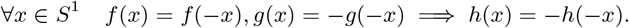

*To see this, let us assume, without loss of generality, that the function f has odd symmetry about the origin, and g has even symmetry about the origin. Fourier series coefficients of an odd function is also odd, implying f*_n_ = *−f*_*−*n_, *while those of an even function is even, implying g*_n_ = *g−n. Previously in Equation (72), we found Fourier series coefficients of the function h to be h*_n_ = 2*πf*_n_*g*_*−*n_. *Taken together, these coefficients are odd h*_n_ = *−h*_*−*n_ *and proves that the function h is also odd*.Δ

##### Remark 8.

*Let f* : *S*^2^ *→* ℝ *be a bounded continuous function. Integral of f* (*x − y*_0_, *x − z*_0_) *over x can be identified with another function h that depends on the difference y*_0_ *− z*_0_ *∈ S*^1^ *as follows:*

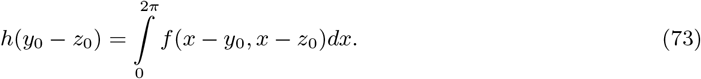

*To see this, let us expand f into its 2D Fourier series representation*

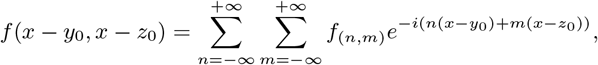

*where f*_(n,m)_ *denotes the Fourier series coefficient for the n*^th^ *and m*^th^ *harmonic over the first and second arguments of f, respectively. Using this expansion, we can compute the integral of f as follows:*

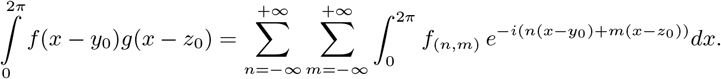

*When m≠ −n, the integral vanishes. Thus, we can reduce the double summation to a single summation with m* = *−n and simplify the integral as follows:*

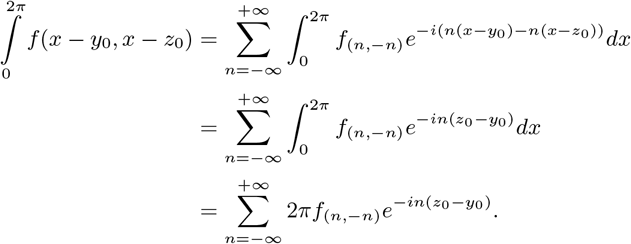

*The resulting summation can be identified as the Fourier series expansion of another function, say h* : ℋ^1^ *→* ℝ, *such that*

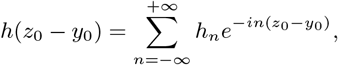

*where h*_n_ := 2*πf*_(n,*−*n)_ *denotes the Fourier series coefficients of the function h*. Δ

### 6.3 Control Theory Reveals an Algorithmic Stability Criteria of Gain Recalibration

#### 6.3.1 Derivation of a reduced-order model for the gain update rule

Here, we derive a reduced-order model for the gain update rule that determines the temporal changes in the PI gain. As will be evident below, the model can capture a broad class of neural mechanisms, including but not limited to activity dependent plasticity like Hebbian plasticity.

We begin our derivation by expressing Equation (59), an analytical expression of the PI gain, in a more compact form, namely,

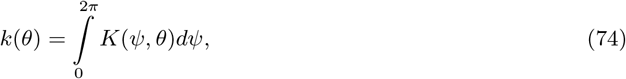

where *K* denotes a function that quantifies the contribution of each location *ψ* in the neural space to the PI gain for a given position representation *θ* as follows:

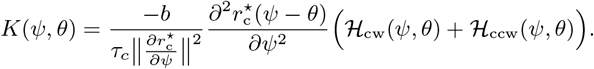

In the remainder of this section, we refer to *K* as the distributed gain function.

Gain recalibration requires the distributed gain function *K* to vary over time. To address this requirement, we extend the domain of *K* to depend explicitly on time, i.e., *K*(*t, ψ, θ*). We are interested in a function, termed the “gain update rule”, that relates the temporal change in *K* across the entire neural space to a change in average PI gain. To find this relation, we first take the time-derivative of the spatial average of Equation (74), which yields

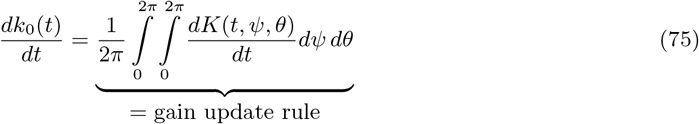

We then search for an explicit, closed-form solution to this integral, assuming that the temporal changes in *K* are driven by two factors:

1. Activity dependent factors: We consider two distinct forms of it:
  a. As the first type of activity-dependent plasticity, we consider that the function *K* at each location *ψ* is modified locally by the firing rates of a finite number of neurons at a specific location relative to *ψ*. An example would be Hebbian plasticity of the synapses between two neurons at the same location *ψ* or at another location a certain distance away from *ψ*. What variables are needed to model the firing rates? Recalling the ansatz solutions to the firing rates of neurons in the central and rotation rings in Equations (27) and (29), we know that firing rate of any neuron in the rotation and central rings can be computed using three variables : *ψ − θ, ψ*, and *v*. As for the visual ring and velocity neurons, we assume that *ψ − θ*^⋆^ and *v* are sufficient to compute firing rate of any neuron, similar to the ansatz solutions in Equations (7) and (25). Therefore, four variables are sufficient to compute the firing rate of any neuron in the ring attractor network: i)*ψ − θ*, ii)*ψ − θ*^⋆^, iii)*v*, iv)*ψ*. Using these variables, we can model the local activity-dependent changes in the distributed gain function *K* via

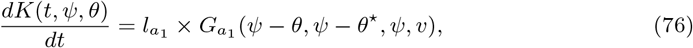

where 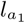 denotes the learning rate, and 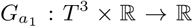 denotes the learning rule. Note that, although we determined the arguments of the function 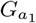 based on the calculation of the firing rates, we do not enforce a specific form for it to make our model as general as possible rather than restricting it to a certain learning rule like Hebbian plasticity between two specific neurons. This way, the learning rule 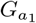 can imitate a broad class of neural mechanisms, including (but not limited to) firing rate dependent plasticity like Hebbian plasticity.
  b. As the second type of activity dependent plasticity, we consider an extended version of the first type. In this extended case, the distributed gain function *K* at neural location *ψ* is changed both locally by the firing rates of a finite number of neurons at a particular location and globally by all the other neurons’ firing rates across the entire neural space. Two motivating examples can be described as follows: 1) Hebbian plasticity in the recurrent connections of the central ring 2) Plasticity mediated by neuromodulators whose release is controlled based on population activity. Recall that the four variables, namely, i)*ψ − θ*, ii)*ψ − θ*^⋆^, iii)*v*, iv)*ψ*, are sufficient to compute the firing rate of any neuron in the network. Since we now consider plasticity that depends on both local activity and global activity, we can model the activity-dependent changes in the distributed gain function *K* at *ψ* via an integral of a function that depends on the same four variables, from which the local firing rates can be computed, as well as additional three variables, i)*ϕ− θ*, ii)*ϕ− θ*^⋆^, iii)*ϕ*, from which the firing rate of other neurons can be computed. This integral is over *ϕ* and takes the form

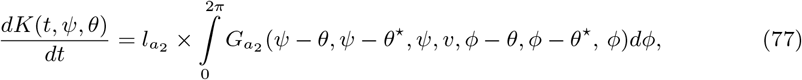

where 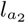 denotes the learning rate, and 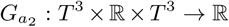 denotes the learning rule. It is again worth recalling that the learning rule 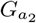 can capture a broad class of neural mechanisms since we do not enforce a particular functional form for it.
2. History dependent effects: To make our model even more general, we take possible history dependent effects into account. However, to limit the mathematical possibilities of how these effects can act, we consider a specific form inspired from the synaptic weight decay well known in the context of Hebbian plasticity [82]. Thus, we model history dependency as a linear decay of the distributed gain function *K* toward a preconfigured value as follows:

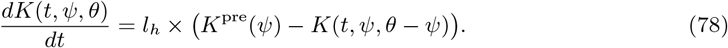

Here, *l*_h_ denotes the forgetting rate of *K* decay toward its preconfigured value *K*^pre^ : *S*^1^ *→* ℝ defined at each location *ψ* in the neural space.

Combining Equations (76), (77), and (78) via superposition, we obtain a general model for the gain update rule in Equation (75), namely,

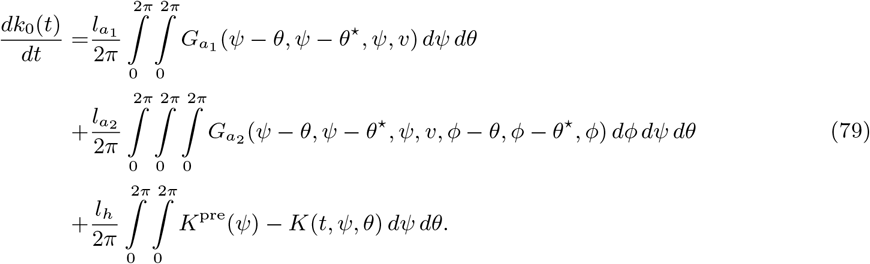

This equation shows that the gain update rule within the ring attractor admits a complicated analytical model. However, as we prove in the next proposition (Prop. 9), this complicated model can be equivalently reduced to a much simpler form. Proof of the proposition makes use of the Remark 8 presented at the end of the previous section.

##### Remark 9.

*The gain update rule in Equation (79) can be equivalently written as*

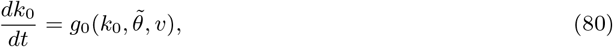

*where g*_0_ : ℝ *×*ℝ *× S*^1^ *→* ℝ *denotes a function that depends on the current gain k*_0_, *the animal’s velocity v, and the positional error* 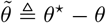 *in the ring attractor’s representation relative to the visual drive’s representation. To see this, we can analyze each line in Equation (79) and show that the integral in each line admits a closed-form, simpler expression as a solution:*

*Line 1. By substituting* 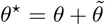 *(note that* 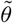 *denotes the error as in the main text) and changing the order of integration, we can rewrite the double integral in the first line as*

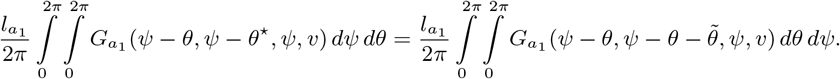

*According to Remark 8, the inner integral reduces to a function that depends on three terms:* 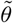, *as the difference between the first and second arguments of* 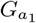, *ψ as the third argument of* 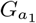, *and v as the last argument of* 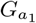 . *The outer integral further reduces to a function that depends on two terms* 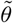 *and v by marginalizing over ψ. These reductions can be expressed mathematically as follows:*

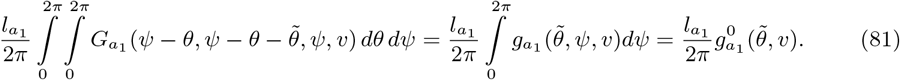

*Here*, 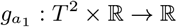 *denotes the function that describes the result of the inner integral, and* 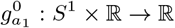 *denotes the function that describes the final result of the double integral*.

*Line 2. Performing the inner most integration over ϕ, we can reduce the triple integral in the second line to a double integral, namely*,

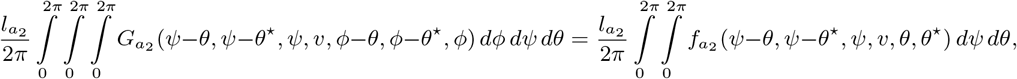

*where* 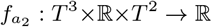 *denotes a function that describes the result of the inner most integral. Since the last two arguments of* 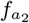 *can be trivially computed from its first three arguments, we can omit the first two arguments of* 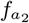, *which simplifies the above equation to*

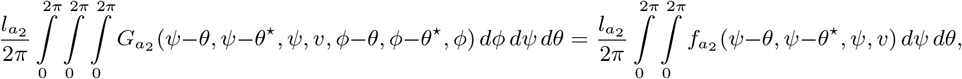

*We then follow the same steps as Line 1. Briefly, we substitute* 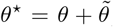, *change the order of integration in the double integral, and use Remark 8 to show that there exists a function* 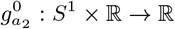 *such that*

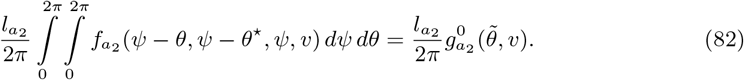

*Line 3. Let us denote the integral of K*^*pre*^ : *S*^1^ *→* ℝ, *the PI gain’s preconfigured value, by* 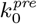, *namely*,

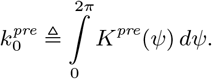

*Taking the spatial average of Equation (74) over θ, we can compute k*_0_ *as*

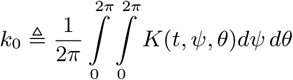

*Using these equations, we can simplify the double integral in third line as follows:*

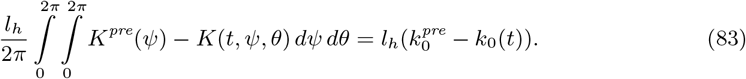

*Combining these simplifications, we obtain a much simpler, closed-form analytical expression for the gain update rule in Equation (79)* :

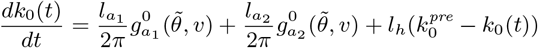

*To make the right hand-side even more general rather than enforcing the above form, we can represent it with a function, say g*_0_, *that depends on* 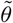, *v, and k*_0_. *Consequently, we find that the gain update rule in Equation (79) reduces to*

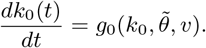

Δ

#### 6.3.2 Proof of Proposition 1 (Necessary conditions for complete gain recalibration)

Here, we provide a Proposition to formally prove Equation (13), which identifies the necessary condition for convergence of the error coordinates 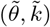 to the origin. For this proof, we analyze the error dynamics in Equation (12), reproduced below:

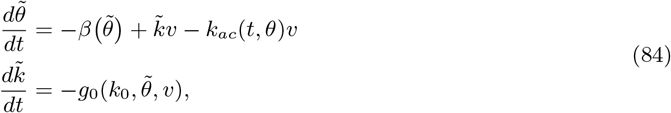

where 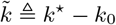. Moreover, we employ the following remark and assumption for the proof.

##### Remark 10.

*A nonzero velocity v≠* 0 *is necessary for stable recalibration, i*.*e*., *if the animal is stationary (i*.*e*., *∀t, v*(*t*) = 0*), then k*_0_ *does not stably converge to k*^⋆^.

*This necessity can be easily verified with a counter example. For v* = 0, *and using the fact that β has odd symmetry about the origin as previously shown in Section 6.2.5, it is easy to show that* 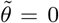 *and* 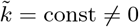, *is a (constant) solution to the error dynamics in Equation (12), and thus* 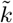 *does not converge to* 0 *(in fact, it does not even change). Intuitively, this is because without moving, the animal can never get any information about its gain*. Δ

##### Remark 11.

*Let B*_*δ*_(0) *⊂* ℝ^2^ *be an open ball of radius δ centered at the origin. If* 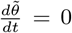 *for all* 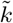 *when* 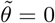*and either one of the following two conditions are satisfied, then the origin* 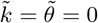 *is not asymptotically stable:*

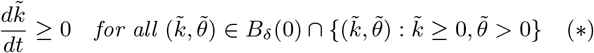

*or*

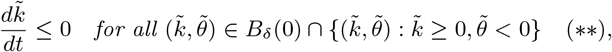

*This result can be verified by considering trajectories beginning in the ball B*_*δ*_(0) *⊂* ℝ^2^ *in* 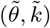 *coordinates. Without loss of generality, let us assume the first condition (labeled with* (*\**)*) given above is true. Asmyptotic stability of the origin requires any point in the ball B*_*δ*_(0) *⊂* ℝ^2^ *to converge to the origin. For trajectories beginning inside the first quadrant*, 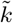 *is positive and constrained to increase further because of* 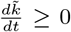. *To avoid* 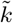 *increasing indefinitely, trajectories must cross the horizontal or vertical axis and go into a different quadrant. However, because* 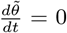 *on the vertical axis and* 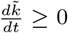 *on the horizontal axis, trajectories are confined to the first quadrant, where* 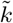 *increases monotonically. Thus, the origin cannot be asymptotically stable*. Δ

##### Assumption 5.

*The AC magnitude, k*_ac_, *of the PI gain function is upper bounded by the distance between the PI gain function’s current average (denoted by k*_0_(*t*)*) and its average at the steady state (denoted by* 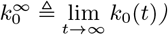, *namely*,

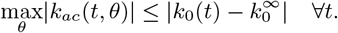

*This implies, k*_*ac*_ (*t, θ*) = 0, *when* 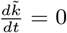. Δ

##### Remark 12.

*The stabilizing feedback β* : *S*^1^ *→* ℝ *is an odd function that crosses zero at the origin (i*.*e*., *β*(0) = 0*) with a positive slope (i*.*e*.,*β*^*′*^(0) *>* 0*). Since this function is odd periodic, it attains a symmetric maximum and minimum, namely*,

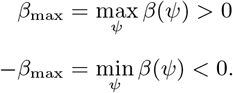

##### Remark 13.

*Let* 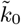 *and v*_0_*≠* 0 *be constants that satisfy* 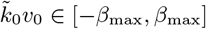. *If* 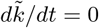 *at* 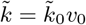, *then k*_*ac*_ = 0 *by Assumption (5), and there exists a solution to* 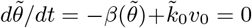 *at* 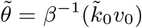. *This implies* 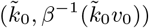 *is an equilibrium point*. Δ

##### Proposition 2

(Necessary conditions for complete gain recalibration). *Let 𝒦 be a bounded open interval in* ℝ. *If the PI gain’s spatial average converges to the visual gain (i*.*e*., 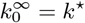*) for all k*^⋆^ *in 𝒦 and for all nonzero velocities of the animal (i*.*e*., *∀t, v*(*t*)*≠* 0*), then it requires*

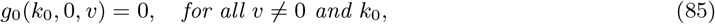

*as well as the sign requirement*

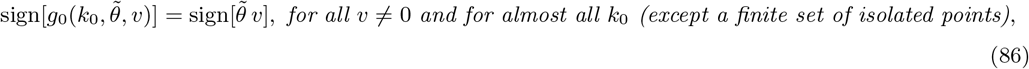

*in a non-empty open interval of the positional error* 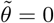 *other than* 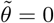.

**Proof of Equation (85)**. We first make an observation: If *k*_0_ converges to *k*^⋆^, then 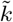 and *k*_ac_ converge to zero, resulting in exponentially stable 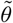 dynamics, i.e., 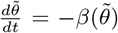, because of *β*^*′*^(0) *>* 0. Thus, convergence of *k*_0_ to *k*^⋆^ *∈ D* implies that the origin 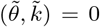 is asymptotically stable for any *k*^⋆^ *∈ D*.

Based on this observation, we can prove the proposition by showing that *Equations (85) and (86) are necessary for asymptotic stability of the origin* 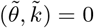 *for any k*^⋆^ *∈ D*.

We first prove the necessity of Equation (85) for the asymptotic stability of the origin. This equation describes the behavior of *g*_0_ at 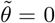. Since the stability of the origin 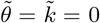 implies that the origin is an equilibrium, namely that the error dynamics in Equation (84) must satisfy

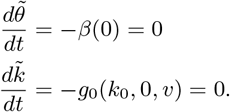

The first equation is automatically satisfied since *β* function has odd symmetry about the origin. The second equation, however, requires

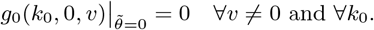

Therefore, gain update rule *g*_0_ satisfies the requirement (85) in the statement of the proposition. □

**Proof of Equation (86)**. We prove the necessity of Equation (86) for asymptotic stability of the origin through contradiction, namely, by showing the contrapositive statement:

*The origin* 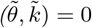 *is not asymptotically stable if there exists a constant v* = *v*_0_*≠* 0 *and a set 𝒦 of k*_0_*’s that is infinite or includes a limit point* 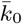*such that*

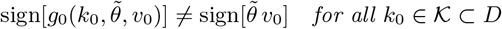

*in any non-empty open interval of*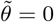 .

Recalling that, by Bolzano-Weierstrass theorem, every bounded infinite set in ℝ has at least one limit point, we find it sufficient to prove this contrapositive statement for a set *𝒦* of *k*_0_’s that has a limit point 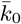. Note that the limit point 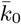 is not necessarily included in *𝒦*, but, according to the definition of a limit point, in every neighborhood of 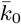, there exists a point in *𝒦* other than 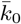.

For each 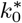 in *𝒦*, the following is true based on the italicized contrapositive statement above:

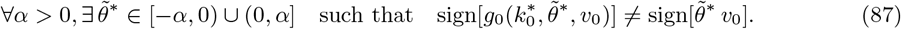

Without loss of generality, let us assume *v*_0_ *>* 0. Under this assumption, the inequality in (87) implies either one of the following, mutually exclusive cases:

- For some 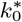 in *𝒦*, there exists a point in every open interval of 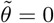 other than 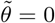, where *g*_0_ is zero, namely,

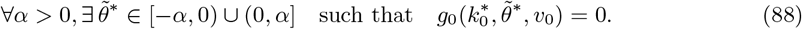

This implies that the zero set of the restriction of the function *g*_0_ to 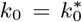 and *v* = *v*_0_, i.e., 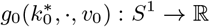, has a limit point at 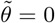. By analyticity of *g*_0_, we infer that this restriction is a zero function, namely,

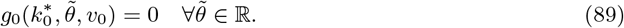
- If Equation (88) is not true for some 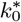 in *𝒦*, then there exists an open interval of 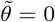 where *g*_0_ is not zero anywhere but 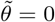. Taken together with Equation (87), this implies

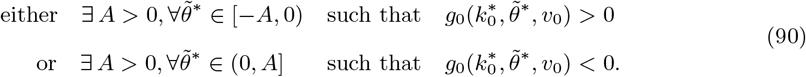

Let us define

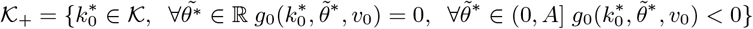

as the set of points in *𝒦* that satisfy either Equation (89) or the second line in Equation (90). Adopting a similar notation, we can define the set of points in *𝒦* that satisfy either Equation (89) or the first line in Equation (90) as

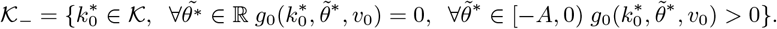

Note that the union of these sets cover the entire *𝒦*, namely, *𝒦*_+_ *∪ 𝒦*_*−*_ = *𝒦*. Moreover, a limit point 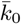 of the set *𝒦* is also a limit point of *𝒦*_+_ or *𝒦*_*−*_. To see this, suppose 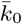 is not a limit point *𝒦*_*−*_. Thus, there exists *ϵ*^⋆^ *>* 0 such that the interval 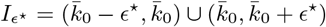 does not include any point in *𝒦*_*−*_, i.e., *I*_ϵ_ *∩ 𝒦*_*−*_ = *∅*. Since 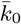 is a limit point of *𝒦* and *𝒦* = *𝒦*_+_ *∪ 𝒦*_*−*_, for every *ϵ* that is 0 *< ϵ < ϵ*^⋆^, the interval *I*_ϵ_ includes an element of 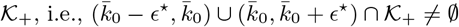, implying that 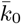 is a limit point of *𝒦*_+_.

Without loss of generality, let us assume 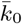 is a limit point of the set *𝒦*_+_. Recall that a limit point of a set is not necessarily included in the set. We show that, regardless of whether the limit point 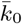 is included in *𝒦*_+_ or not, the origin 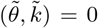 is not asymptotically stable for some visual gain *k*^⋆^ *∈ D*. To this end, consider an open ball 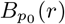 centered at 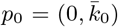 with radius of radius *r < β*_max_. For a given *r*, pick *δ*_r_ *>* 0 that solves 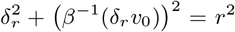 (the solution exists since *r < β*_max_ ). This *δ*_r_ ensures that the interval 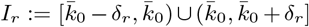 is contained in the ball 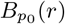. For every *r >* 0, this non-empty interval exists. Since 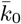 is a limit point of *𝒦* _+_, there exists a point 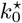 in *I*_*r*_ *∩ 𝒦* _+_ that satisfies either Equation (89) or the second row of Equation (90) for every *r >* 0. Below, we investigate each case and show that the origin 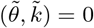 is not asymptotically stable for some *k*^⋆^ *∈ D*.

Case 1. Consider first that 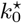 satisfies Equation (89). Let us choose the visual gain 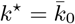. Equation (89) implies that the change in the gain error 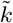 is zero, namely,

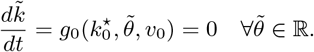

When this equation is satisfied, *k*_ac_ = 0 by Assumption 5. It further follows from Remark 13 that the point 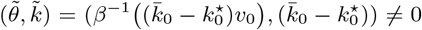 is an equilibrium. Thus, the origin cannot be asymptotically stable as there exists another equilibrium point other than the origin in any open ball of radius *r >* 0.

Case 2. Consider next that, for some *r*^⋆^ *>* 0, there is no 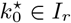 satisfying Equation (89), i.e., Case 1 is not true. Thus, for all *r ∈* (0, *r*^⋆^], there exists 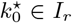 satisfying the second row of Equation (90) such that

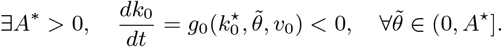

On the other hand, depending on whether the limit point 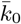 is inside the set *𝒦* _+_ or not, the change in the gain *k*_0_ can be positive or negative, namely, there exists *Ā >* 0 such that

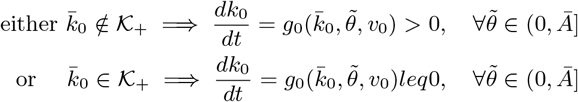

Subcase 1. In the subcase of 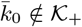, continuity of *g*_0_ implies that there exists an intermediate point 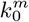 in the open interval connecting 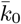 and 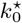 such that

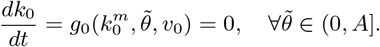

where *A* = min(*A*^⋆^, *Ā*). Analyticity of *g*_0_ further requires the function to be zero for all values of 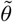 at 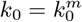, namely,

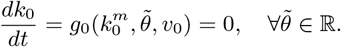

This raises a contradiction, as we assumed that Case 1 is not true. Thus, this subcase is not possible.

Subcase 2. Consider the remaining subcase 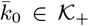. For *v* = *v*_0_, this case implies the function *g*_0_ is nonpositive everywhere in a half-closed region, namely, 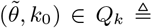, (0, min(*A*^⋆^, *Ā*)] *× I*_*k*_ where *I*_*k*_ is the open interval connecting 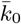 and 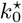 . Choose a visual gain in this open interval, i.e., *k*^⋆^ *∈ I*_*k*_. Since 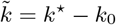, this implies

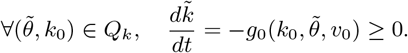

One can find an open ball *B*_*δ\**_ (0) that is entirely contained in *Q*_*k*_ in the right-half plane. Since 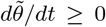 in the right-half of this ball and 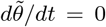 on the vertical axis 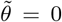 by Equation (85), we can employ Remark 11 to conclude the origin is not asymptotically stable.

Therefore, gain update rule *g*_0_ satisfies the requirement (86) in the statement of the proposition. □

#### 6.3.3 Extension of Proposition 1 to Partial Recalibration

##### Corollary 1

*Let the animal’s velocity be constant v* = *v*_0_*≠* 0 *and K be a bounded open interval in* ℝ. *If the PI gain’s spatial average converges to* 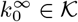 *for a given k*^⋆^, *then the following statements are true*.

1. *There is an an equilibrium point* 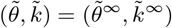 *that satisfies*

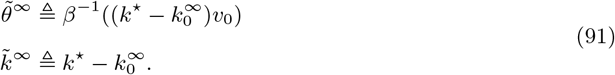
2. *Assume* 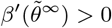. *The gain update rule g*_0_ *satisfies*

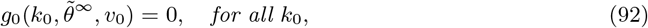

*as well as the sign requirement*

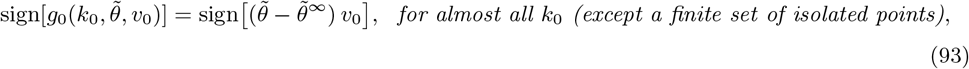

*in a non-empty open interval of the positional error* 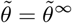 *other than* 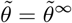.

**Proof of Statement 1**. Convergence of *k*_0_ to 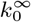 implies that 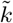 converge to

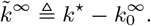

After convergence, 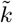 dynamics satisfy

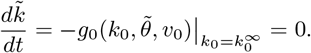

On this nullcline of 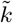, the PI gain’s AC component *k*_*ac*_ is zero by Assumption 5. Substituting *k*_*ac*_ = 0 into 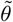 dynamics in (84), we obtain 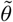 dynamics on the nullcline of 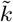 as

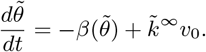

Assuming 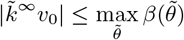, we can solve 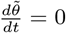 for 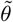, namely,

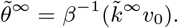

Therefore, 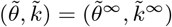 is an equilibrium point of the dynamics in Equation (84). □

**Proof of Statement 2**. Before proving this statement, we first perform the change of coordinates 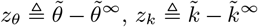. Under this coordinate change, the point 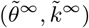 in 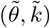 coordinates maps to the origin (0, 0) in (*z*_*θ*_, *z*_*k*_) coordinates. By Statement 1, the origin (*z*_*θ*_, *z*_*k*_) = 0 is an equilibrium point. Also, note that 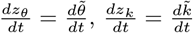 by the chain rule.

In these new coordinates, wee can reformulate the dynamics in Equation (84) as

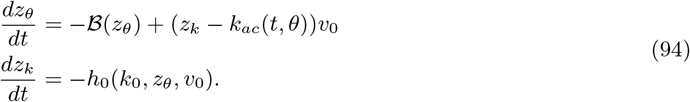

where

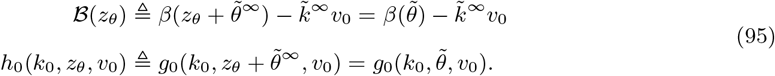

Recall that the PI gain’s AC component *k*_*ac*_ in this equation satisfy the inequality in Assumption 5. By adding subtracting *k*^⋆^ to the right-hand side of that inequality, we can reformulate it in the new coordinates as

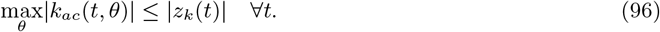

Furthermore, definition of *ℬ* and the fact 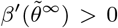 imply that the function *ℬ* satisfy *ℬ*(0) = 0 and *ℬ*^*′*^(0) *>* 0, the same properties possessed by the function *β* (i.e., *β*(0) = 0 and *β*^*′*^(0) *>* 0).

Based on these reformulations, we can rephrase the second statement in the corollary as follows: *Let 𝒦*_*z*_ *be a bounded open interval in* ℝ *and ℬ*^*′*^(0) *>* 0. *If z*_*k*_ *converges to the origin for any* 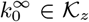, *then the function h*_0_ *satisfies*

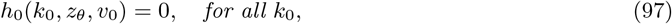

*as well as the sign requirement*

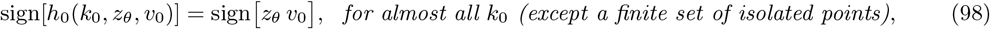

*in a non-empty open interval of z*_*θ*_ = 0 *other than z*_*θ*_ = 0.

Note that (*z*_*θ*_, *z*_*k*_) dynamics in Equation (94) are isomorphic to 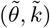 dynamics in Equation (84). Moreover, the rephrased statement is a special case of the statement in Prop. 2 for *v* = *v*_0_*≠* 0 and 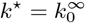. Therefore, using Prop. 2, we prove the above rephrased statement of the corollary. □

#### 6.3.4 Proof of Proposition 2 (Sufficient Conditions for Gain Recalibration)

Here, we provide a Proposition to formally prove Equation (15) in the main text, which identifies the sufficient condition for convergence of the error coordinates 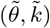 to the origin. Throughout the proof, we make use of the following remarks.

##### Remark 14.

*As stated previously in Equation (96), the term k*_*ac*_ *is bounded by the absolute value of z*_*k*_. *We can model this relation more directly by including z*_*k*_ *into the arguments of the function k*_*ac*_, *namely k*_*ac*_(*t, θ, z*_*k*_). *△*

##### Remark 15.

*Suppose visual gain k*^⋆^ *and the animal’s velocity v*_0_ *are given. In this case, θ*^⋆^ = *k*^⋆^*v*_0_*t is determined completely by the time t. Since one can compute* 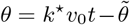, *the gain’s spatial variation can be modeled, without loss of any generality, as a function that depends on* 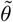 *instead of θ, i*.*e*., 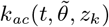.

*Moreover, since the gain’s spatial variation is zero-mean and periodic over θ*^⋆^ *for a given* 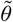, *the function* 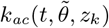 *is zero-mean and periodic over time t with period T* = 2*π/*(*k*^⋆^*v*_0_), *namely*,

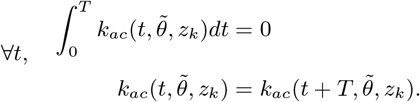

Δ

*Finally, since* 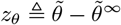, *the second argument of this function can be considered as a variable that depends on z*_*θ*_ *for a fixed* 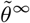 *instead of* 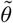, *namely, k*_*ac*_(*t, z*_*θ*_, *z*_*k*_).Δ

##### Proposition 3

(Sufficient conditions for gain recalibration). *For a given velocity v* = *v*_0_*≠* 0 *and visual gain k*^⋆^, *if the gain update rule satisfies the slope condition*

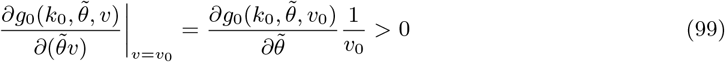

*everywhere in the solution set of* 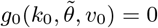, *then PI gain k*_0_ *converges a steady-state value*.

*Proof*. Let us assume, without loss of generality, that the steady-state value of *k*_0_ is 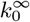, corresponding to 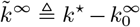 as the steady-state value of 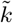. A necessary condition for convergence to these steady-state values were previously given in Corollary 1. According to this corollary, the function *g*_0_ must satisfy

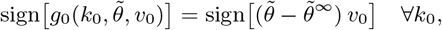

where 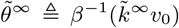 denotes the steady-state value of 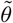. This requirement implies that 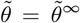 constitutes the solution set of 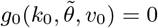 for all *k*_0_, namely,

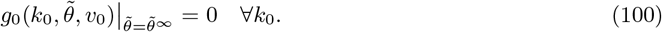

With this in mind, we now proceed with an investigation of Equation (99), describing the slope condition, to prove its sufficiency for asymptotic stability of the point 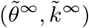. For this investigation, we take a multi-step approach.

1. We first perform a change of coordinates 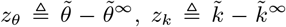, which leads to the same non-autonomous system dynamics as in Equation (94). This change of variables maps the point 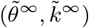, representing steady-state values in the 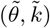 coordinates, to the origin in the (*z*_*θ*_, *z*_*k*_) coordinates. Under this change of coordinates, *k*_0_, the first argument of the function *h*_0_, can be expressed as 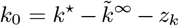. Because 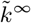 is fixed as a coordinate of the equilibrium point and because visual gain *k*^⋆^ is given, we can re-organize the arguments of the function *h*_0_ without loss of generality as *h*_0_(*z*_*k*_, *z*_*θ*_, *v*_0_). In this new form, the slope condition in Equation (99) can be written as

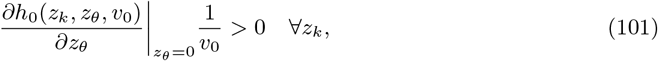

where we used 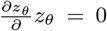 identifies the solution set of *h*_0_(*z*_*k*_, *z*_*θ*_, *v*_0_) = 0 in the new form (see Equation (100) for the respective solution in the old form). This solution set implies

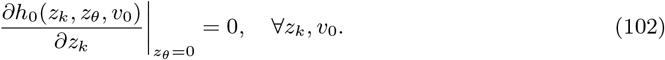
2. In the next step, we work with the term *k*_*ac*_, which denotes the AC component of the PI gain. Recall that, by Equation (96), when *z*_*k*_ = 0, we have *k*_*ac*_ = 0 for all *t, z*_*θ*_, which implies

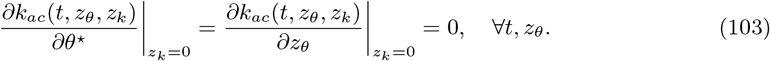
3. Finally, on the basis of our findings in the previous steps, we will show that the system dynamics in Equation (94) is stable as claimed in the proposition. To this end, we first modify Equation (94) by replacing the function *g*_0_ with the function *h*_0_ and by updating the functional form of *k*_*ac*_ as described in Remark 15, which yields

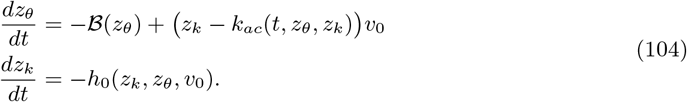

For the stability assessment, we employ the Averaging Theorem from dynamical systems theory that exploits the fact that *k*_*ac*_ is a periodic function over time with zero-mean and period *T* = 2*π/*(*k*^⋆^*v*_0_). According to this theorem, if the autonomous system without the periodic perturbation *k*_*ac*_ is exponentially stable at the origin, then Equation (104), the nonautonomous system with periodic perturbation *k*_*ac*_, attains an exponentially stable periodic orbit around the origin. Moreover, as stated in [83], that premise makes the origin an exponentially stable equilibrium point of the perturbed, nonautonomous system in Equation (104) due to 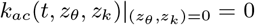. Therefore, to prove the proposition, it is sufficient to show that the origin is an exponentially stable equilibrium point of the autonomous system without the periodic perturbation *k*_*ac*_ when Equation (101) is true. (See Theorem 10.3 in [83] and Theorem 4.1.1 in [84] for more formal, detailed statements of the Averaging Theorem.) For this specific task, we first ignore the *k*_*ac*_ in Equation (104), which leads to the autonomous system without any periodic perturbation, namely

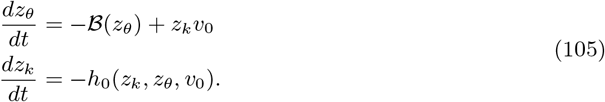

To assess stability of this autonomous system, we linearize it about the origin (*z*_*θ*_, *z*_*k*_) = 0, leading to the following linear time-invariant system:

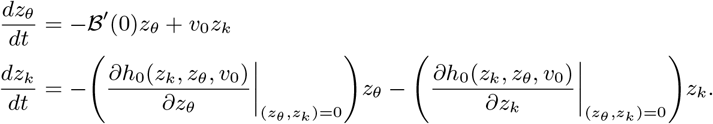

Substituting Equation (102) into this system, we obtain

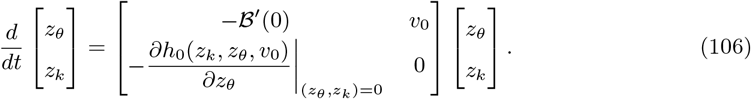

Both eigenvalues of this system are negative, namely,

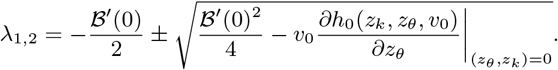

The inequality in Equation (101) ensures that these eigenvalues are negative, i.e., *λ*_1,2_ *<* 0, rendering the origin an exponentially stable equilibrium point of the linear system in Equation (106). Since this linear system was obtained by linearizing Equation (105) about the origin, it follows from Hartman-Grobman theorem [83] that the origin is an exponentially stable equilibrium point of the autonomous system in Equation (105). Moreover, by using the aferomentioned Averaging theorem, we can conclude that the origin is also an exponentially stable equilibrium point of the nonautonomous system in Equation (104). This proves that the PI gain *k*_0_ converges to a steady-state value, namely 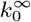, as claimed in the proposition. □

### 6.4 Mechanistic Constraints Reveal Instrumental Role of Positional Error Codes

In this section, we analyze analytical expressions of the average PI gain *k*_0_ to identify the mechanistic constraints for the gain recalibration. We derive two alternative expressions by averaging Equations (59) and (61) over *θ*, which yields

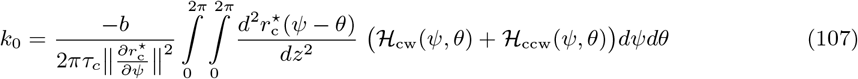

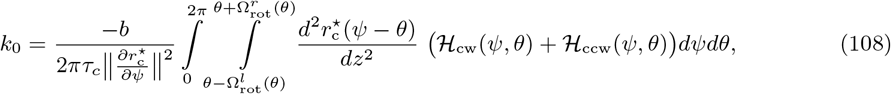

respectively. Also recall that *H*_cw_, *H*_ccw_ the terms must satisfy the symmetry constraint in Remark 3. For the expressions above, this implies

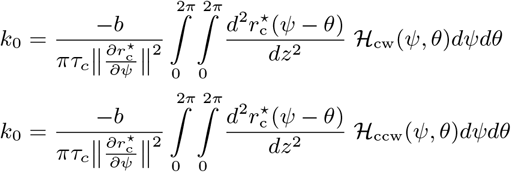

and

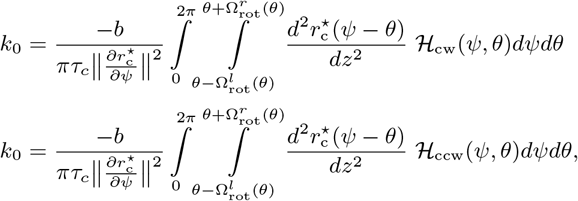

respectively. Using the closed-form analytical expressions of the terms *H*_*cw*_, *H*_*ccw*_ in Equation (46), we can rewrite these expressions more explicitly as

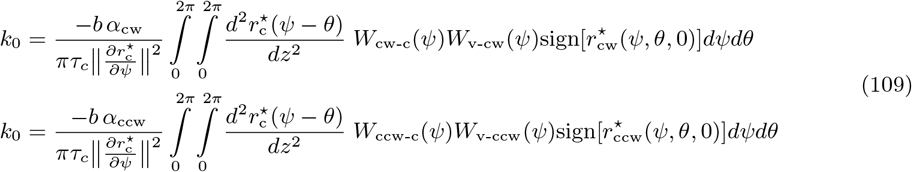

and

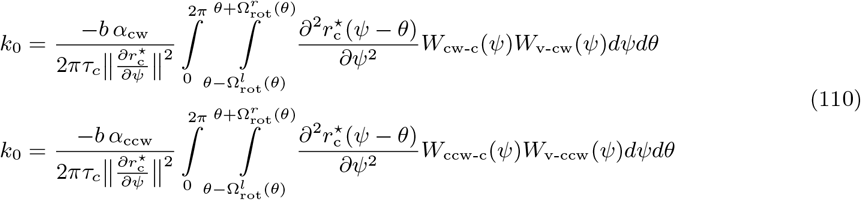

#### 6.4.1 Temporal changes in the synaptic weights of connections between the velocity neurons and the rotation rings

In this subsection, we examine how gain recalibration can be driven by temporal changes in the functions *W*_v-cw_ and *W*_v-ccw_, describing the synaptic weight of connections between the velocity neurons and the rotation rings. To consider the temporal effects of these functions on *k*_0_, we extend Equation (109) to be time-dependent as given below:

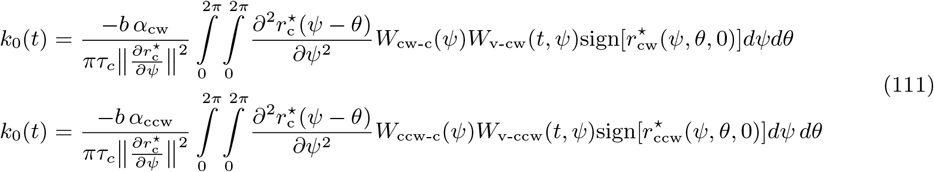

On the other hand, we assume that the remaining parameters of our model are hardwired (i.e., time-invariant), initialized with values inspired by the classical ring attractor models such that

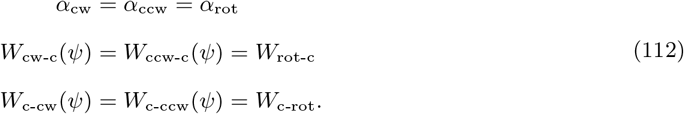

Since the integrands in Equation (111) are finite and attain their extreme values over a compact space, their double integrals in these equations lead to finite constants. This lets us change the order of integration. Doing so and re-arranging the terms, we obtain

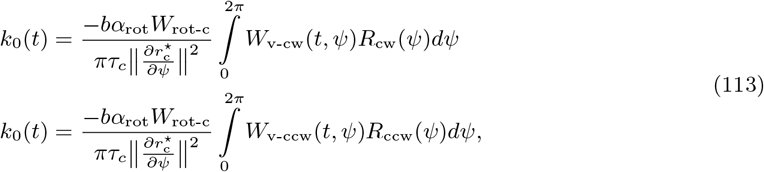

where the functional terms *R*_cw_ and *R*_ccw_ denote

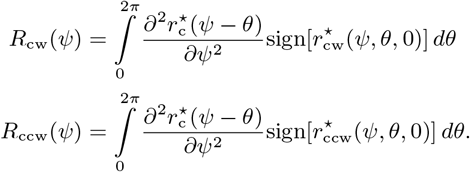

Notice that, by virtue of Assumption 2, the integrands in these terms are negative inside the compact support intervals of 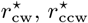 and zero elsewhere. Thus, the functional terms *R*_cw_, *R*_ccw_ are negative.

We now employ the first mean value theorem for definite integrals. According to this theorem, since the functions *R*_cw_, *R*_v-ccw_ are negative everywhere (i.e., not changing sign), there exists constants 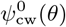, 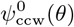 for each *θ* such that Equation (113) reduces to

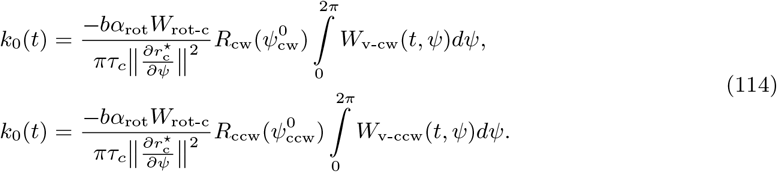

Differentiating this equation with respect to time, we obtain the relation between the temporal change in the synaptic weights and the temporal change in the average gain, namely,

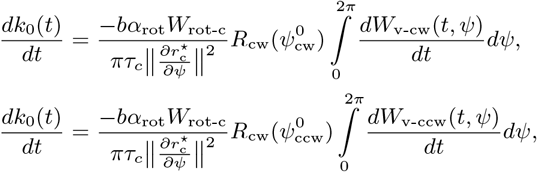

Since the leading coefficient of the integral in each line is positive, Equation (86) that describes the necessary condition for gain recalibration, translates to

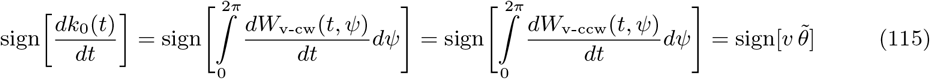

Assuming Hebbian plasticity, we can transform this requirement on the synaptic weights into a requirement on the firing rates of the rotation rings and velocity neurons, namely,

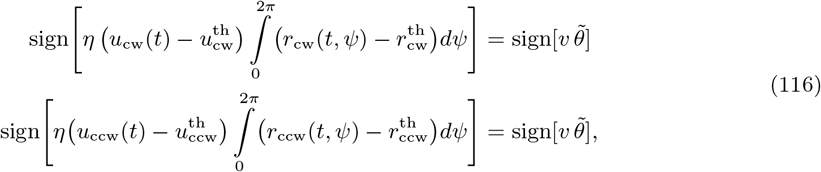

where *η >* 0 denotes the learning rate, and the terms with the superscript *th* denotes the threshold activity-levels. These thresholds must be chosen properly to halt the plasticity when either *v* or 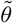 is zero. Thus, we choose the activity thresholds of velocity neurons as follows:

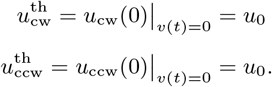

According to the model of velocity neurons in Equation (25), this implies

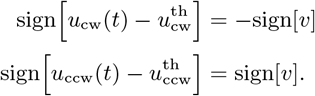

Using this implication and the fact *η >* 0, we can refine the sign requirement in Equation (116) to

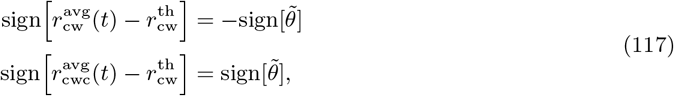

where 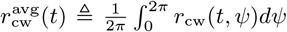 and 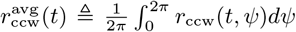 denote the average firing rate of CW and CCW rotation rings, respectively. The sign requirement in this equation implies that these average firing rates encode the positional error 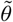 via

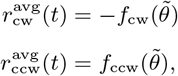

where *f*_cw_ : *U*_cw_ → ℝ, *f*_ccw_ : *U*_ccw_ : R denote strictly increasing functions in the open neighborhoods *U*_cw_, *U*_ccw_ of the origin.

#### 6.4.2 Temporal changes in the synaptic weights of connections between the rotation rings and the central ring

In this subsection, we examine how gain recalibration can be driven by temporal changes in the functions *W*_cw-c_ and *W*_ccw-c_, describing the synaptic weight of connections between the rotation rings and the central ring. To consider the temporal effects of these functions on *k*_0_, we extend Equation (109) to be time-dependent as given below:

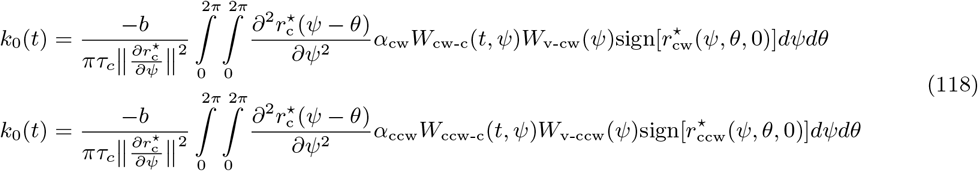

On the other hand, we assume that the remaining parameters of our model are hardwired (i.e., time-invariant), initialized with values inspired by the classical ring attractor models such that

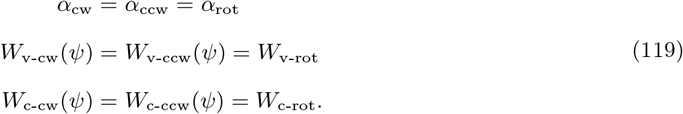

We now apply the steps between Equation (113) and (115), but this time keeping the now plastic weights *W*_cw-c_, *W*_ccw-c_ inside the integral and instead moving the now hard-wired weights *W*_v-cw_(*ψ*) = *W*_v-ccw_(*ψ*) = *W*_v-rot_ outside. This yields the stability requirement of gain recalibration for the temporal changes in the plastic weights *W*_cw-c_, *W*_ccw-c_, namely,

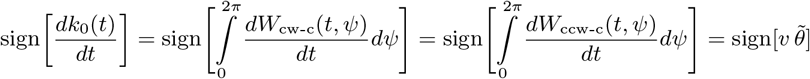

We next assume Hebbian plasticity as the mechanism of these temporal changes in *W*_cw-c_, *W*_ccw-c_. This assumption lets us transform this sign requirement on the synaptic weights into a requirement on the firing rates of the central ring and rotation ring neurons, namely,

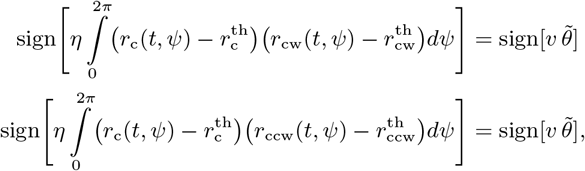

where *η >* 0 denotes the learning rate, and the terms with the superscript *th* denote the threshold activity-levels. Using the fact *η >* 0 and the superscript avg as a shorthand notation for the average firing rates, we can simplify this equation to

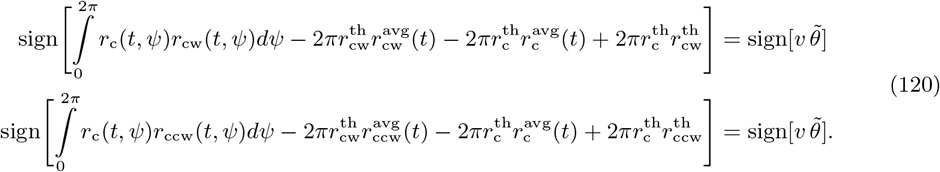

Recall that firing rates *r*_c_,*r*_cw_, and *r*_ccw_ are positive. This lets us employ the first-mean value theorem for the definite integral in Equation (120). According to this theorem, there exists constants 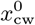 and 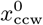 such that

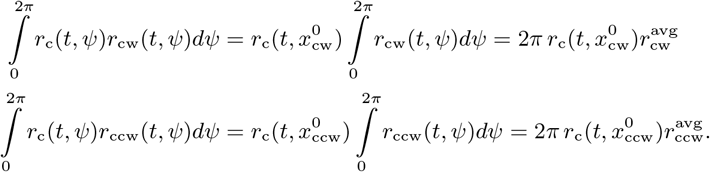

Under the assumption that the shape of the central ring’s activity bump, denoted by 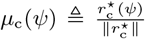 remains constant thanks to attractor dynamics, there exists a constant gain factor 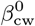 for a given 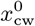 such that 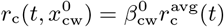. Using this, we can rewrite the above equation as

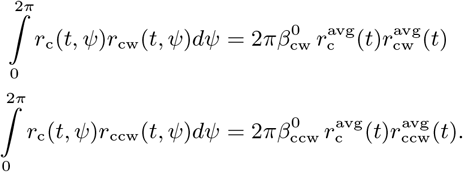

Substituting this equation into (120), we obtain the following simplified sign requirement:

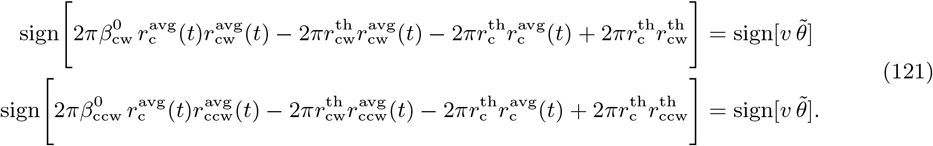

Satisfying each line this equation at a given time *t* and velocity *v* requires that the central ring’s average firing rate 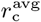 or the rotation rings’ average firing rates 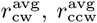 must encode the positional error 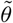 via strictly monotonic changes. More formally, there exists strictly monotonic functions *f*_c_ : *U*_c_ → ℝ, *f*_cw_ : *U*_cw_ → ℝ, *f*_ccw_ : *U*_ccw_ → ℝ in non-empty open neighborhoods *U*_c_,*U*_cw_,*U*_ccw_ of the origin such that at least one of the

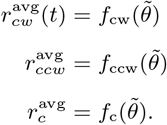

#### 6.4.3 Temporal changes in the tuning slope of velocity neurons

In this subsection, we examine how gain recalibration can be driven by temporal changes in the terms *α*_cw_ and *α*_ccw_, denoting the slope of the CW and CCW velocity neuron’s tuning curves, respectively. To consider the temporal effects of these terms on *k*_0_, we extend Equation (109) to be time-dependent and re-arrange terms, which yields

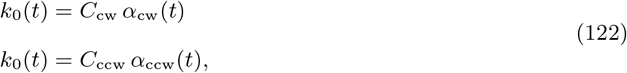

where the positive constants *C*_cw_, *C*_ccw_ take the form

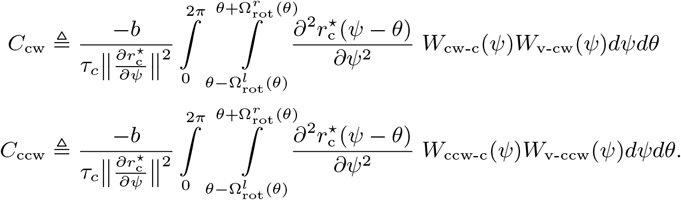

On the other hand, we assume that the remaining parameters of our model are hardwired (i.e., time-invariant), initialized with values inspired by the classical ring attractor models such that

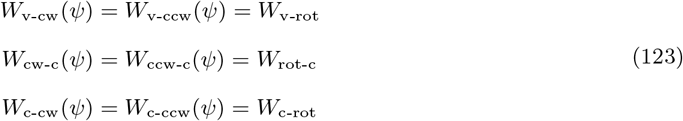

Note that, under these assumptions, the left- and right-widths of rotation rings’ activity bumps during immobility become equal and independent of *θ*, namely,

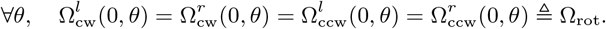

Differentiating Equation (122) with respect to time and using Equation (86), describing the necessary condition for gain recalibration, we obtain the requirements that must be satisfied by the temporal changes in *α*_cw_, *α*_ccw_ for gain recalibration as follows:

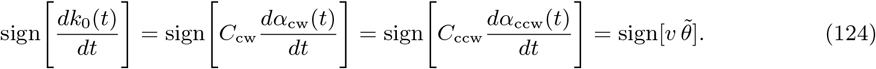

Since *C*_cw_ and *C*_ccw_ are positive, this simplifies to

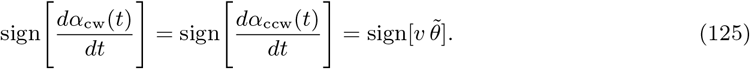

In previous sections, we examined the neural mechanisms of plasticity that can satisfy similar equalities through temporal changes in the synaptic weights. However, for the velocity neurons, our model does not include details to examine the neural mechanisms governing the temporal changes in the slopes of their tuning curves. Alternatively, we can garner insight into the consequences of these temporal changes for the rest of the neurons in our model, assuming that the above equalities are met. In the remainder of this section, we examine how average firing rate of rotation rings are affected by the temporal changes in the slopes of the velocity neurons.

### Implications of Equation (124) on the average firing rate of rotation rings

It can be trivially seen from Equation (23) that the average firing rate of rotation rings varies proportionally with the slopes *α*_cw_,*α*_ccw_. Specifically,

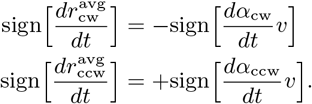

Plugging Equation (125) into the right-hand side, we can further refine this requirement :

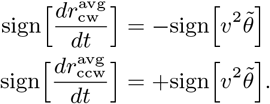

According to these equations, for a given *v*, the temporal change in the rotation rings’ average firing rates must be in the same direction as 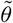, namely,

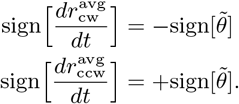

This implies that temporal changes in the rotation rings’ average firing rates can be identified with strictly monotone functions of 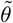 in some neighborhood of the origin. More specifically, we can find two strictly increasing functions *f*_cw_ : *U*_cw_ → ℝ, *f*_ccw_ : *U*_ccw_ → ℝ in the open neighborhoods *U*_cw_, *U*_ccw_ of the origin such that

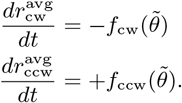

What can we tell about the average firing rates 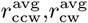 when their temporal changes are strictly monotone functions of 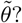 To answer this question, we restrict ourselves to simple 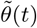 trajectories that do not change sign by crossing zero. Under this condition, one can show that average firing rates of the CW and CCW rotation rings encode the time-integral 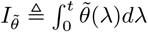 of the positional error via

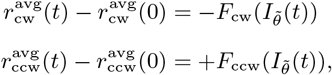

where *F*_cw_ : *𝒰*_cw_ → ℝ, *F*_ccw_ : *𝒰*_ccw_ → ℝ denote strictly increasing functions of 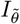 in some non-empty neighborhoods *𝒰*_cw_, *𝒰*_ccw_ of the origin.

#### 6.4.4 Temporal changes in the width of the rotation ring’s activity bump

In this subsection, we examine how gain recalibration can be driven by temporal changes in the terms 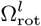 and 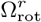, respectively, denoting the left- and right-width of the rotation rings’ activity bumps during immobility. As a first step toward this examination, we extend Equation (108), the analytical expression of the average gain *k*_0_, to be time-dependent as given below:

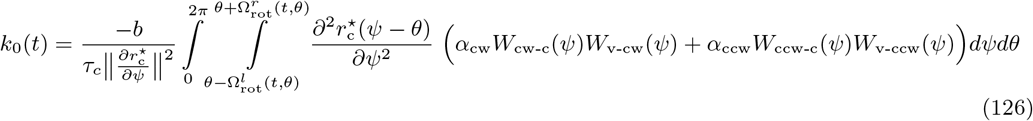

On the other hand, we assume that the remaining parameters of our model are hardwired (i.e., time-invariant), initialized with values inspired by the classical ring attractor models such that

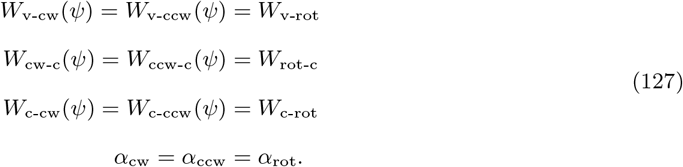

Under these assumptions, 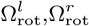 become independent of *θ*, implying that they can change only over time, i.e. 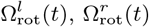. Taking into account this implication and the assumptions in Equation (127), we can simplify Equation (126) to

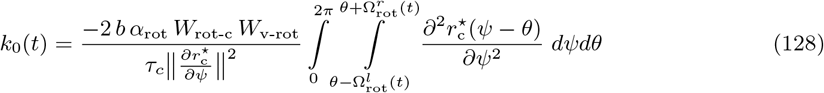

If the widths 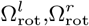 are time-invariant, *k*_0_ in Equation (128) is a constant, implying that the temporal change in *θ* does not lead to any change in *k*_0_. Therefore, taking time-derivative of Equation (128) and ignoring the changes due to *θ*, we obtain the relationship between the temporal change in *k*_0_ and that in the widths 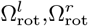 as follows:

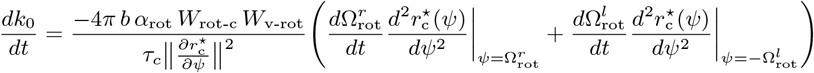

Note that, when deriving this equation, we applied the Leibniz’s integral rule since the widths 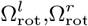 appear in the bounds of an integral in Equation (128).

For gain recalibration, this temporal change in *k*_0_ must satisfy Equation (86), which implies

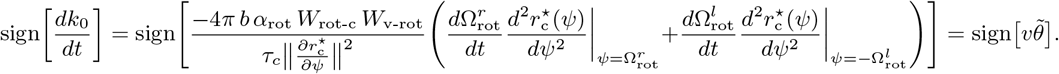

Since the terms 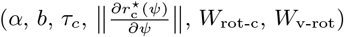 are positive, we can simplify this requirement to

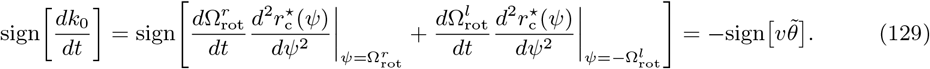

In addition to this equation, the widths 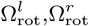 must satisfy the symmetry constraint in Remark 3. Using Equations (46), (127) and applying the steps between Equation (128) and (129), we can express this symmetry constraint in a form that explicitly depends on 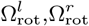 as follows:

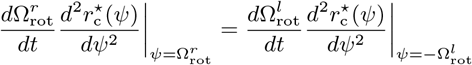

Combining this symmetry constraint with Equation (129), we find that the necessary condition for gain recalibration requires the temporal changes in 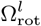 and 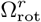 to change according to

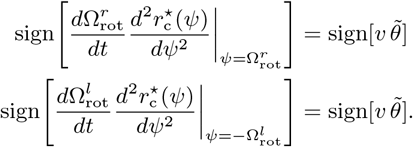

Recall that, by Assumptions 1 and 2, the second derivative 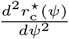 takes a negative value at the points 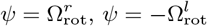. Thus, the above equation can be simplified to

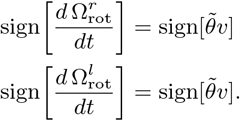

According to these equations, for a given *v*, the temporal change in the rotation rings’ widths must be in the same direction as 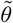, namely,

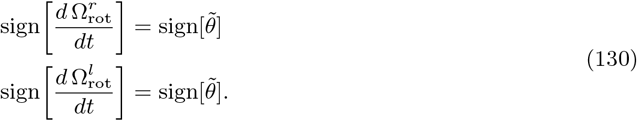

This implies that there exists strictly increasing functions 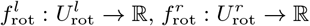 in the open neighborhoods 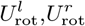 of the origin such that

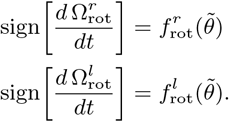

What can we tell about the left- and right-widths 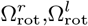 when their temporal changes are strictly increasing functions of 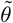? To answer this question, we restrict ourselves to simple 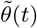 trajectories that do not change sign by crossing zero. Under this condition, one can show that left- and right-widths of the rotation rings encode the time-integral 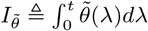 of the positional error via

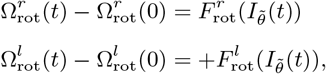

where 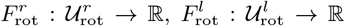 denote strictly increasing functions of 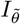 in some non-empty neighborhoods 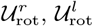 of the origin.

#### 6.4.5 Temporal changes in the norm of the central ring’s persistent activity bump

In this subsection, we examine how gain recalibration can be driven by temporal changes in the term 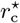, describing the central ring’s persistent activity bump. Specifically, we assume that the activity bump’s shape 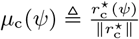 remains invariant while the temporal changes in the norm 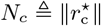 drive the gain recalibration. On the other hand, we assume that the remaining parameters of our model are hardwired (i.e., time-invariant), initialized with values inspired by the classical ring attractor models such that

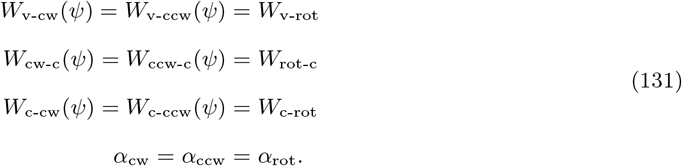

An implication of these assumptions is that 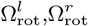 are independent of *θ*. Re-organizing Equation (108), the analytical expression of *k*_0_, in accordance with these conditions, we obtain

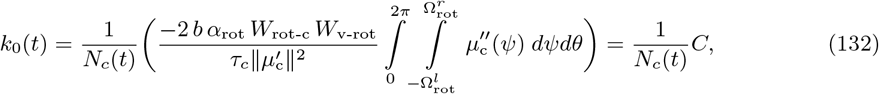

where *C* denotes a positive constant, and *N*_*c*_(*t*) denotes the time-varying norm of the persistent activity bump 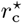 as the driver of the gain recalibration. Also note that, while reorganizing to obtain this equation, we used the following facts

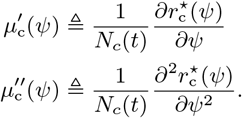

In order to find the relation between the temporal change in *N*_*c*_ and the resulting change in *k*_0_, we simply differentiate Equation (132) with respect to time, which yields,

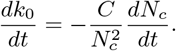

For gain recalibration, this temporal change in *k*_0_ must satisfy Equation (86). This implies

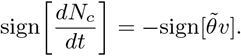

According to this equation, for a given *v*, the temporal change in the norm of the central ring’s activity bump must be in the opposite direction of 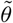, namely,

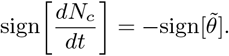

This implies that there exists a strictly increasing function *f*_c_ : *U*_c_ → ℝ in the open neighborhood *U*_c_ of the origin such that

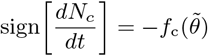

What can we tell about the norm *N*_*c*_ when its temporal changes is a strictly decreasing function of 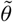? To answer this question, we restrict ourselves to simple 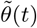 trajectories that do not change sign by crossing zero. Under this condition, one can show that the norm *N*_*c*_ of the central ring’s activity bump encode the time-integral 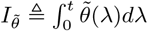 of the positional error via

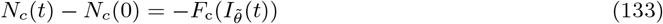

where *F*_c_ : *𝒰*_c_ → ℝ denotes a strictly increasing function of 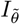 in some non-empty neighborhood *U*_c_of the origin.

Note that the norm *N*_*c*_ is related to the average firing rate 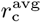 via

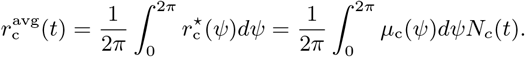

Since the integral 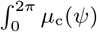 is constant (i.e., time-invariant), Equation (133) implies that the average firing rate 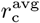 of the central ring encodes the time-integral of 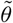 via similar monotonic changes.

### 6.5 Analytical proof of gain recalibration in the proposed ring attractor model

The proposed model employs plasticity in the connections between the velocity neurons and the rotation ring neurons. Assuming Hebbian plasticity, we can model this plasticity through

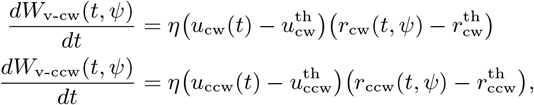

where *η >* 0 denotes the learning rate, and the terms with the superscript *th* denotes the threshold activity-levels. As we previously derived in Appendix 6.4.1, both synaptic weights *W*_v-cw_ and *W*_v-ccw_ are related to the average gain *k*_0_ via Equation (114). Differentiating each line in this equation, averaging over these time-derivatives of the lines, and then using the model of Hebbian plasticity (given above), we obtain an equation for how firing rates of velocity and rotation rings neurons drive the gain recalibration in our proposed attractor model, namely,

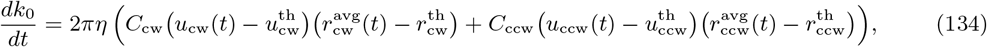

where the terms *C*_cw_, *C*_ccw_ denote the positive constants

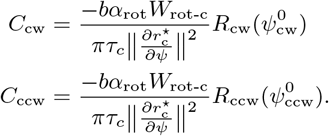

Recall that, firing rate of velocity neurons encode the velocity *v* via

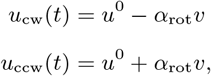

where *α*_rot_ denotes the absolute value of the tuning curves’ slopes. Moreover, the rotation rings encode *v* and error 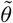 conjunctively. This conjunctive coding takes the form of a superposition, which can be approximated as a linear model, namely,

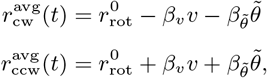

where *β*_v_ and 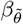 denote absolute value of slopes due to *v* and 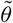, respectively. Using these firing-rate equations and the threshold choices

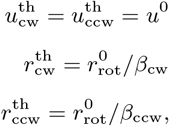

we can reduce Equation (134) to a gain update rule *g*_0_ that is a function of 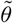 and *v*, namely,

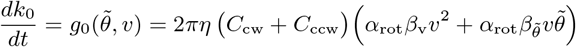

For a given *v* ≠ 0, this gain update rule satisfies

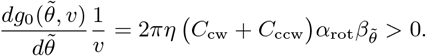

As per Prop. 3, this inequality guarantees that the proposed model achieves stable gain recalibration.

## Acknowledgments

This work was supported by National Institutes of Health grants R01 NS102537 (N.J.C., J.J.K.), U01 NS131438 (N.J.C.), and R01 MH118926 (J.J.K., N.J.C.). We thank Ravikrishnan Jayakumar, Kathryn Hedrick, Bharath Krishnan, Manu Madhav, Francesco Savelli, and Kechen Zhang.

## Footnotes

1 We follow the convention *F*_x-y_ to denote a directional quantity *F*, such as a synaptic weight or current, from neurons of population *x* to neurons of population *y*.

2 Let *f, g* : *S*^1^ *→* ℝ. 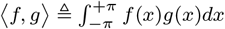 denote the inner product, and 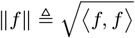 denote the *L*_2_ norm. Projection operator is defined as 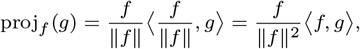.

